# Flat-to-curved transition during clathrin-mediated endocytosis correlates with a change in clathrin-adaptor ratio and is regulated by membrane tension

**DOI:** 10.1101/162024

**Authors:** Delia Bucher, Felix Frey, Kem A. Sochacki, Susann Kummer, Jan-Philip Bergeest, William J. Godinez, Hans-Georg Kräusslich, Karl Rohr, Justin W. Taraska, Ulrich S. Schwarz, Steeve Boulant

## Abstract

Although essential for many cellular processes, the sequence of structural and molecular events during clathrin-mediated endocytosis remains elusive. While it was believed that clathrin-coated pits grow with a constant curvature, it was recently suggested that clathrin first assembles to form a flat structure and then bends while maintaining a constant surface area. Here, we combine correlative electron and light microscopy and mathematical modelling to quantify the sequence of ultrastructural rearrangements of the clathrin coat during endocytosis in mammalian cells. We confirm that clathrin-coated structures can initially grow flat and that lattice curvature does not show a direct correlation with clathrin coat assembly. We demonstrate that curvature begins when 70% of the final clathrin content is acquired. We find that this transition is marked by a change in the clathrin to clathrin-adaptor protein AP2 ratio and that membrane tension suppresses this transition. Our results support the model that mammalian cells dynamically regulate the flat-to-curved transition in clathrin-mediated endocytosis by both biochemical and mechanical factors.

## Introduction

Clathrin-mediated endocytosis (CME) is an essential uptake pathway that relocates membrane or extracellular cargo into the cell to regulate multiple cellular functions and cell homeostasis^1^. During CME, the clathrin coat is assembled to form a clathrin-coated pit (CCP) that after dynamin-mediated scission from the plasma membrane (PM) leads to the formation of a clathrin-coated vesicle (CCV)^2^. This process is coordinated by numerous adaptor and accessory proteins^1,3^. Electron microscopy (EM) of clathrin coated structures (CCS) has shown the architectural complexity of the clathrin meshwork organized into hexagons and pentagons^4,5^. From this EM analysis, it was proposed that a CCV initiates as a flat clathrin lattice that is then rearranged to form a curved CCP^4,6,7^. However, for topological reasons this requires a major ultrastructural rearrangement of the clathrin lattice which appeared to be dynamically difficult and energetically costly^8–12^. For these reasons, this notion was replaced by a general belief that CCS grow with a constant curvature (constant curvature model, Fig. 1a)^8,9,13^ and that flat CCS are distinct from CCPs and serve different purposes^14–16^. This model was supported by the finding that purified clathrin triskelia self-assemble into curved clathrin baskets *in vitro*^17,18^. Recently, correlative light and electron microscopy (CLEM) analyses provided experimental evidence that CCS first grow flat to their final size and then acquire curvature (constant area model, Fig. 1a)^19^. However, this study did not measure the dynamics of CCP formation directly, and it did not identify the cellular factors that might determine when the flat-to-curved transition occurs. Thus a comprehensive understanding of the dynamic process of coat rearrangement, of the temporal aspects of flat-to-curved transition and of what governs this ultrastructural rearrangement during CME is still missing.

**Figure 1:**
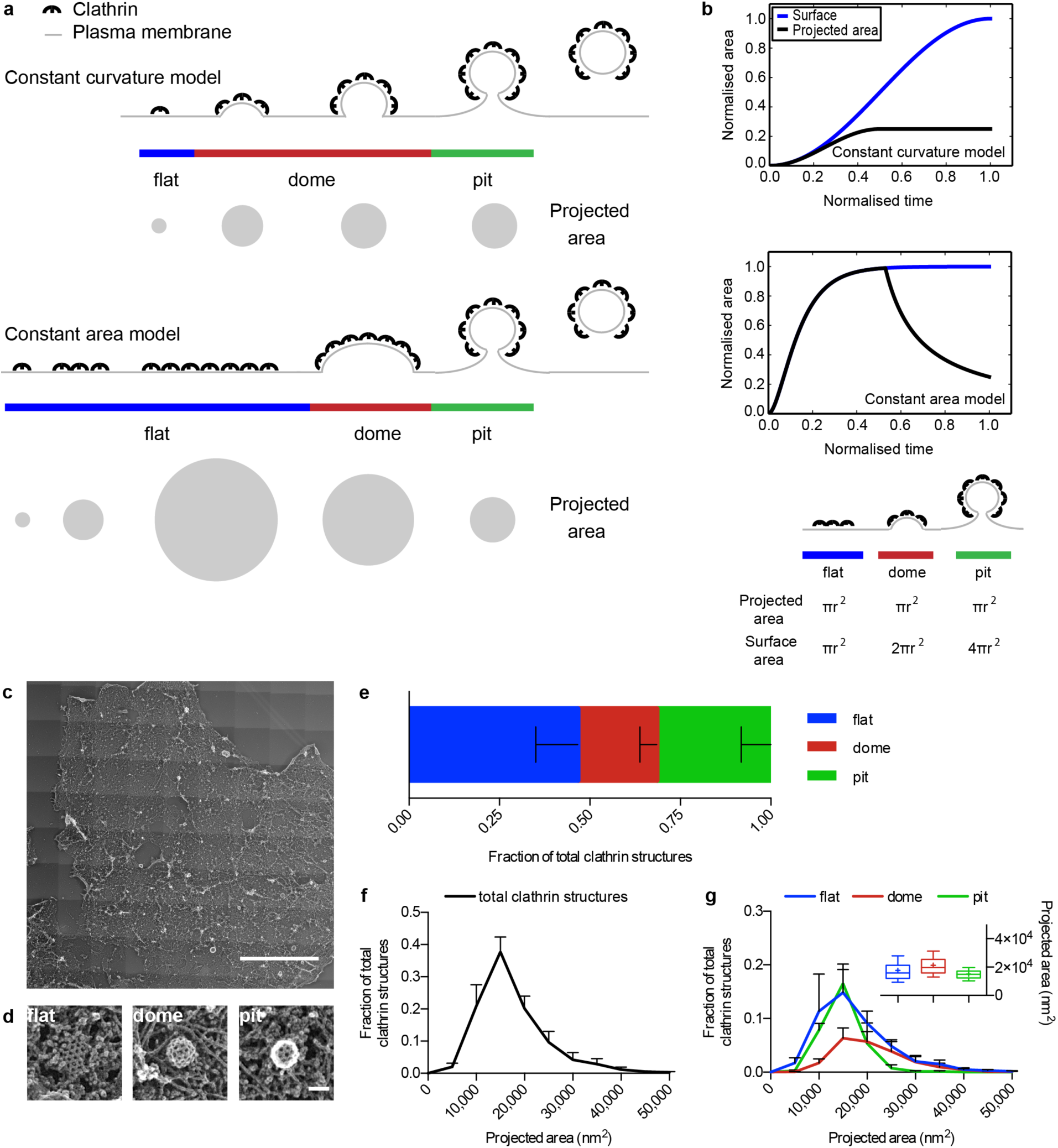
Comprehensive ultrastructural characterisation of CCS in BSC-1 cells by TEM. (a) Schematic of the constant curvature and constant area models. The stages of different curvature (flat (blue), dome (red), pit (green)) and the variation of projected area, which can be assessed during TEM imaging, is depicted for both growth models. (b) Difference between projected area (black) and surface area (blue) during the course of CCP formation according to the two models. The schematic illustrates the relationship between projected area and surface area for flat, dome (approximately a hemisphere), and pit (approximately a complete sphere) CCS. (c) TEM of metal replica from unroofed PM, overview of whole membrane, scale bar: 10μm. (d) Examples of flat, dome, and pit structures, scale bar: 100nm. (e) Fraction of flat (blue), dome (red) and pit (green) CCS in whole PM of BSC-1 cell. (f) Projected area distribution of all CCS (black) measured by TEM. (g) Projected area distribution of the different clathrin morphologies (flat, dome, pit). A box/whiskers plot of the projected area is shown in the inset. Mid-line represents median, cross represents the mean and the whiskers represent the 10 and 90 percentiles. Results are calculated from three different membranes (number of CCS per membrane: 746, 869, and 739); means with SD are shown.

In this work, we combine mathematical modelling of individual endocytic event dynamics and CLEM analysis to provide a comprehensive description of the dynamic ultrastructural rearrangement of the clathrin coat during CME. We demonstrate that CCPs indeed initially grow as flat arrays, but that their reorganisation into curved structures occurs before reaching their full clathrin content. We correlate this flat-to-curved transition with a change in the AP2/clathrin ratio and show that it is governed by biophysical properties of the PM. Our findings provide a unifying view of the dynamic process of coat rearrangement during CME and our approach constitutes a methodological framework to further study the fine-tuned spatio-temporal mechanism regulating coat assembly.

## Results

### EM and CLEM analysis of CCS do not support existing growth models

To address whether CCP formation follows the constant curvature model or the constant area model (Fig. 1a)^13^, we chose BSC-1 cells, a widely used cellular model to study CME^10,14,20^. BSC-1 cells present homogenous CME events in regard to both lifetime as well as intensity profiles and lack the long-live flat clathrin-coated plaques^10,15^ (Supplementary Fig. 1 and Fig. 1c-g). Both models predict different growth profiles for the surface and projected area during CCP formation. The constant curvature model implies that the projected area will quickly be smaller than the surface area. In contrast, the constant area model implies that both projected and surface areas initially show similar growth but then the projected area should drop significantly as bending starts (Fig. 1b).

To comprehensively characterise the ultrastructural organisation of CCS in BSC-1 cells, we performed TEM of metal replicas from unroofed PMs (Fig. 1c). We confirmed that CCS are not altered by the unroofing procedure using stimulated emission depletion (STED) super-resolution microscopy of intact and unroofed cells. The number and size distribution of CCS were indeed similar between intact and unroofed cells (Supplementary Fig. 2). CCS in TEM images of whole PM sheets were counted, categorised as flat, dome or pit structures (Fig. 1d-e) and their size was measured as projected area (Fig. 1a, f, and g). For the constant curvature model, we would expect no flat structures at all and no dome structures that exceed the projected area of pits (Fig. 1a-b). In contrast our EM data reveals that around 50% of the CCS in BSC-1 correspond to flat CCS (Fig. 1e) and that a large fraction of the flat and dome structures have a projected area larger than the projected area of the pits (Fig. 1g). Since BSC-1 cells do not have clathrin-coated plaques^10,15^, these results demonstrate that the constant curvature model cannot explain the CCS size distribution, in agreement with the recent results by Avinoam et al^19^. Although the existence of flat CCS seems to argue in favour for the constant area model, we would expect that some flat structures have the same projected area as the surface area of fully formed pits (Fig. 1a-b). Since the surface area of a spherical pit (4πr^2^, Fig. 1b) is four times larger than its projected area (πr^2^, Fig. 1b), we would expect the mature flat structures to have around four times the projected area of pits (Fig. 1b). Instead, we found no flat structures at all with a projected area four times larger than the mean projected area of CCP (Fig. 1g). Additionally, the constant area model would imply that the projected area of dome structures (which resembles a hemisphere) is reduced by a factor of two when converting to CCP. Instead we found only a slight increase of the mean projected area of domes compared to pits (Fig. 1g, distribution and inset box/whiskers). Together, these observations argue against the constant area model.

To further challenge the two growth models, we used a CLEM approach^21,22^ (Fig. 2a). BSC-1 cells were immunostained with a clathrin heavy chain antibody and the fluorescence intensity of CCS (Fig. 2b) was correlated to their size and ultrastructural organisation measured using TEM of metal replicas (Fig. 2c). For the constant curvature model, we would expect the intensity to increase with increasing contact angle (Fig. 1a-b). In case of the constant area model, we would expect equal intensity for the largest flat, domes, as well as all pit structures (Fig. 1a-b). However, our CLEM analysis clearly revealed that flat and dome structures have similar fluorescence intensities while pits tend to display higher fluorescence intensities (Fig. 2b and d). In conclusion, our TEM and CLEM analyses argue that neither of the proposed growth models fully explains the observed ultrastructural distribution and corresponding fluorescence intensities of CCS in BSC-1 cells.

**Figure 2:**
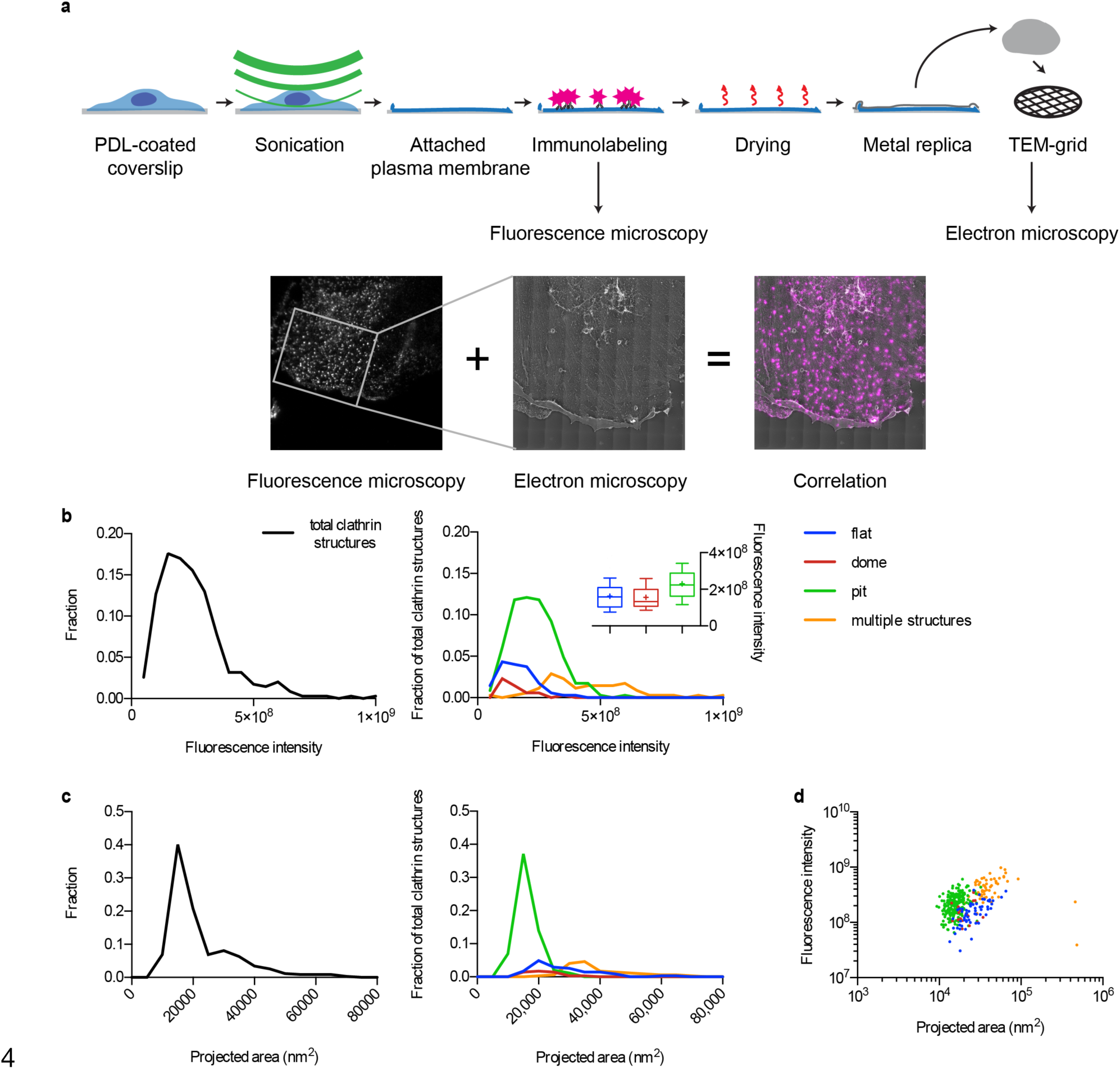
CLEM of CCS. (a) Schematic representation of the CLEM approach (Upper part). Cells growing on poly-D-lysine (PDL)-coated coverslips were unroofed by sonication. Attached PMs were immunostained and imaged using FM. Samples were then critical-point dried and the metal replica was created and lifted from the sample onto a TEM grid for imaging in TEM. (Lower part) The FM and EM pictures were then correlated to combine their information (See methods). Unroofed PMs were immunostained using an anti-clathrin heavy chains antibody. The white inset box represents the area observed by TEM. (b) Fluorescence intensity distribution (clathrin heavy chain antibody, X22) of all CCS (black line, left panel) and of flat (blue), dome (red), pit (green) CCS, and multiple structures which cannot be distinguished by fluorescence microscopy (orange) (right panel). A box/whiskers plot of the fluorescence intensity is shown in the inset. Mid-line represents median, cross represents the mean and the whiskers represent the 10 and 90 percentiles. (c) Projected area distribution of all CCS (black line, left panel) and of the different clathrin morphologies (right panel). (d) Correlation of size and fluorescence intensity of all CCS sorted by their different morphologies. Graphs show one representative CLEM result with a total of 347 CCS from one PM.

### Modelling of CCP assembly reveals bending of the coat before reaching full surface area

Although EM of metal replicas is a high-resolution microscopy technique revealing detailed information about the size and ultrastructure of CCS, it only provides snapshots of the dynamic process of CCP assembly^23^. In contrast, live fluorescence microscopy (FM) of CME mostly allows the characterisation of the dynamics of different proteins during the formation of CCS but does not provide ultrastructural information^9^. To obtain a more comprehensive dynamical picture, we used mathematical modelling of clathrin growth behaviour to combine the ultrastructural information from EM with the dynamic information obtained from total internal reflection fluorescence (TIRF) microscopy of fluorescently tagged clathrin light chain (CLC). As the formation of CCS is a complex process with numerous yet unknown variables, here the modelling approach is used to combine and compare the different data sets (FM, TEM and CLEM), with minimal assumptions on the underlying mechanisms. Since we found a substantial number (around 50%) of flat CCS with projected areas similar to pits (Fig. 1e and g), we rule out the possibility that the constant curvature model might be the dominant growth behaviour in BSC-1 cells. We therefore modelled the growth of flat CCS first as circular planar discs that grow to a finite size before bending (Fig. 3a). Mathematically there is only one type of growth equation that can explain why growth should stop at a finite patch radius, namely association over the edge and dissociation over the area of the patch (Supplementary Information). By fitting this growth equation and assigning the three different morphologies to the individual intensity profiles obtained by TIRF microscopy (Fig. 3b and Supplementary Fig. 1g-i), we could calculate the size and morphology distribution of CCS as predicted by the constant area model (Fig. 3c). Comparison of the calculated distribution to the acquired EM data reveals that the ratio between the flat, dome, and pit structures is biased toward flat structures compared to the EM data (Fig. 3g). Additionally, the means of the predicted size distributions of flat and pit structures are clearly separated with a shift of the flat projected area towards bigger sizes (Fig. 3c and i box/whiskers and Fig. 3h). In agreement with our TEM and CLEM results, our mathematical modelling approach thus demonstrates that the constant area model does not correctly describe the assembly process of CCPs in BSC-1 cells.

**Figure 3:**
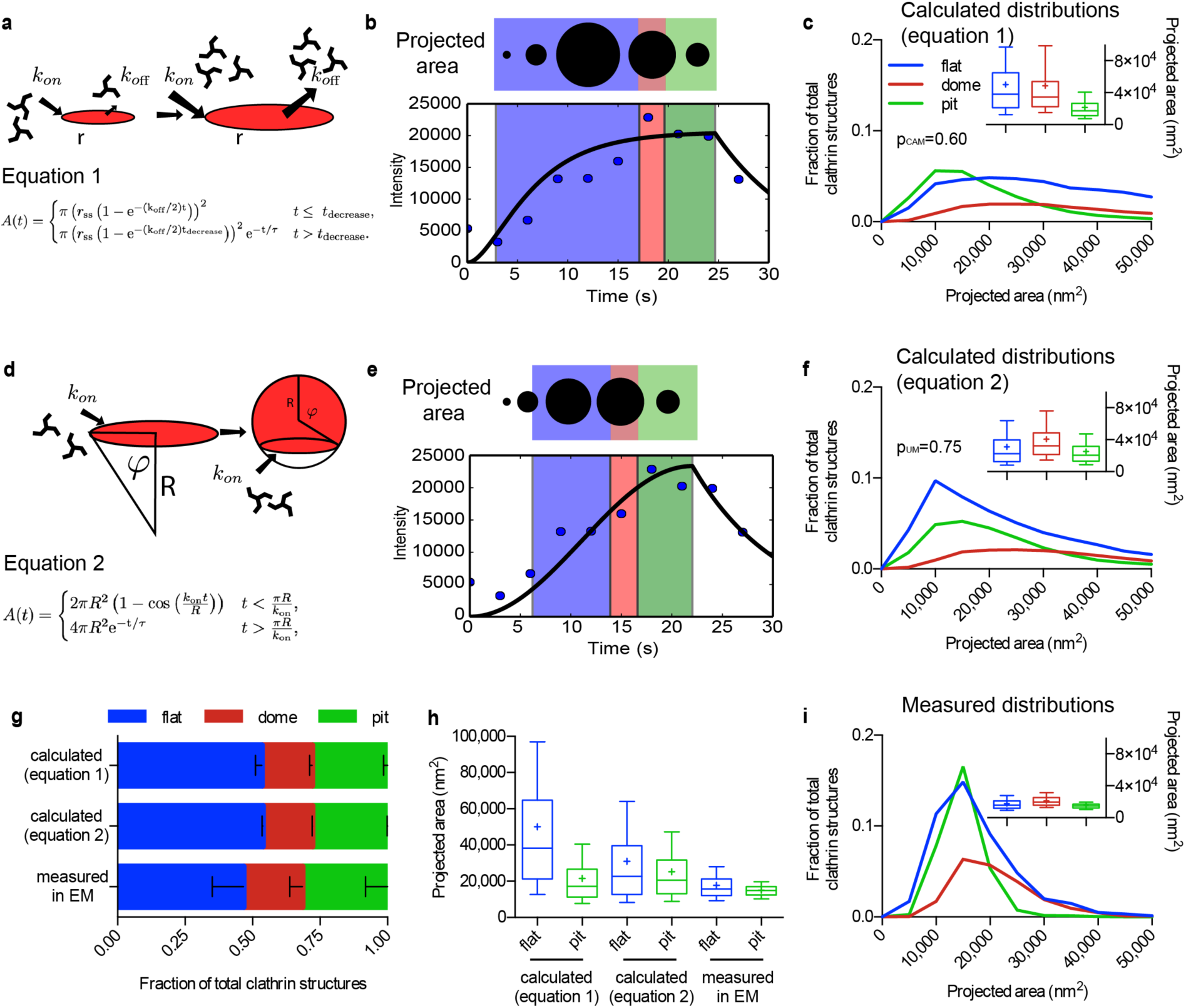
Mathematical modelling of CCS growth from intensity profiles of individual CME events. (a) Mathematical representation of the constant area model, flat-to-curved transition happens at the time when clathrin reaches its final content. (b) Example of a clathrin intensity track fitted by the constant area model. Blue dots represent measured intensity of a single CME event; black line represents the fit with equation 1. The schematic illustrates the calculated size and curvature, flat (blue), dome (red), and pit (green). (c) Calculated projected area and curvature distributions of the CCS according to the constant area model for 4927 FM tracks of 4 different cells. P-value of Welch’s t-test to compare the predicted to the measured distribution in panel h. A box/whiskers plot of the projected area is shown in the inset. Mid-line represents median, cross represents the mean and the whiskers represent the 10 and 90 percentiles. (d) Mathematical representation of the updated growth model where a flat clathrin patch grows and the flat-to-curved transition happens before reaching the final clathrin content. (e and f) same as (b and c) but using equation 2. (g) Comparison of the predicted ratio of flat, dome, and pit structures from both growth model (equation 1 (a) and equation 2 (d)) and the distribution obtained from TEM imaging. Results are calculated for 4927 FM tracks of 4 different cells; means with SD are shown. (h) Direct comparison of the projected area distribution of flat and pit structures calculated by equation 1 and 2 as well as measured in EM, box/whiskers plot. (i) Measured projected area and curvature distributions of the CCS from TEM data as shown in Fig. 1.

Given the high proportion of flat structures (Fig. 1e), we reasoned that CCS start as flat structures and then acquire curvature before reaching the full clathrin content (Fig. 3d and Supplementary Information). In contrast to the patch growth model (Fig. 3a), now the system has an intrinsic mechanism to stop growth, namely formation of a sphere. The patch growth is included as the initial regime. In contrast to the constant curvature model, however, here we do not fix the radius of the sphere. The resulting growth equation was again fitted to the individual intensity profiles of CME events (Fig. 3e) (Supplementary Information). As before, the ratio between the flat, dome, and pit structures was biased towards flat structures (Fig. 3g) but now the calculated size and morphology distribution fit the EM data better than the distribution according to the constant area model. The means of the predicted projected area of both the flat and pit have similar sizes (Fig. 3f and h, box/whiskers). These findings strongly support a model where assembly of a CCP initiates flat and then acquires curvature before it acquires its final clathrin content.

### Starting point of acquiring curvature is marked by a change in the AP2/clathrin ratio

The flat-to-curved transition of a CCS requires major ultrastructural reorganisation of the coat^11^. To acquire curvature, according to Euler’s theorem the hexagonal organisation of the clathrin triskelia needs to reorganise into a polyhedral assembly including 12 pentagons^11^. The clathrin lattice in flat structures is mostly composed of hexagons^4,5^. Although it has been shown using FRAP that the clathrin coat is highly dynamic, which is a prerequisite for such rearrangement^19,24,25^, it is still puzzling what regulates the organisation of triskelia in the coat and what coordinates the flat-to-curved transition. It was proposed before that the ratio of the adaptor AP2 to clathrin changes within the growth of CCP^22,26,27^. Therefore, we correlated the relative amount of AP2 and the ultrastructural organisation of CCS. We performed CLEM analysis using BSC-1 cells expressing AP2 fused to GFP (Fig. 4a) and correlated these results to clathrin immunostaining CLEM (Fig. 4b). To find the relationship between fluorescence intensity and the surface of CCS, the measured projected area needs to be corrected for the curvature to obtain the surface area of the CCS (Fig. 1b). For flat structures, projected area and surface area are identical, thus we used the linear regression of flat coats as a reference to correct the projected area of both domes and pits. Assuming the geometry of a hemisphere for domes and an almost complete sphere for pits we expect a correction factor of ≤2 for domes and 2<x≤4 for pit structures if the relationship between fluorescence intensity and surface area is independent of curvature (Fig. 1b). The correction factors inferred for domes and pits were 1.4 and 2.8, respectively, which are within the expected values (Fig. 4d-e and Fig. 1b). Strikingly, the correction factors for AP2 CLEM were smaller than expected, especially for the pit structures (domes: 1.2; pits: 1.7) (Fig. 4c and e). This reveals that the AP2/clathrin ratio in a CCS differs depending on its curvature and that this ratio is reduced within the coat as curvature increases.

**Figure 4:**
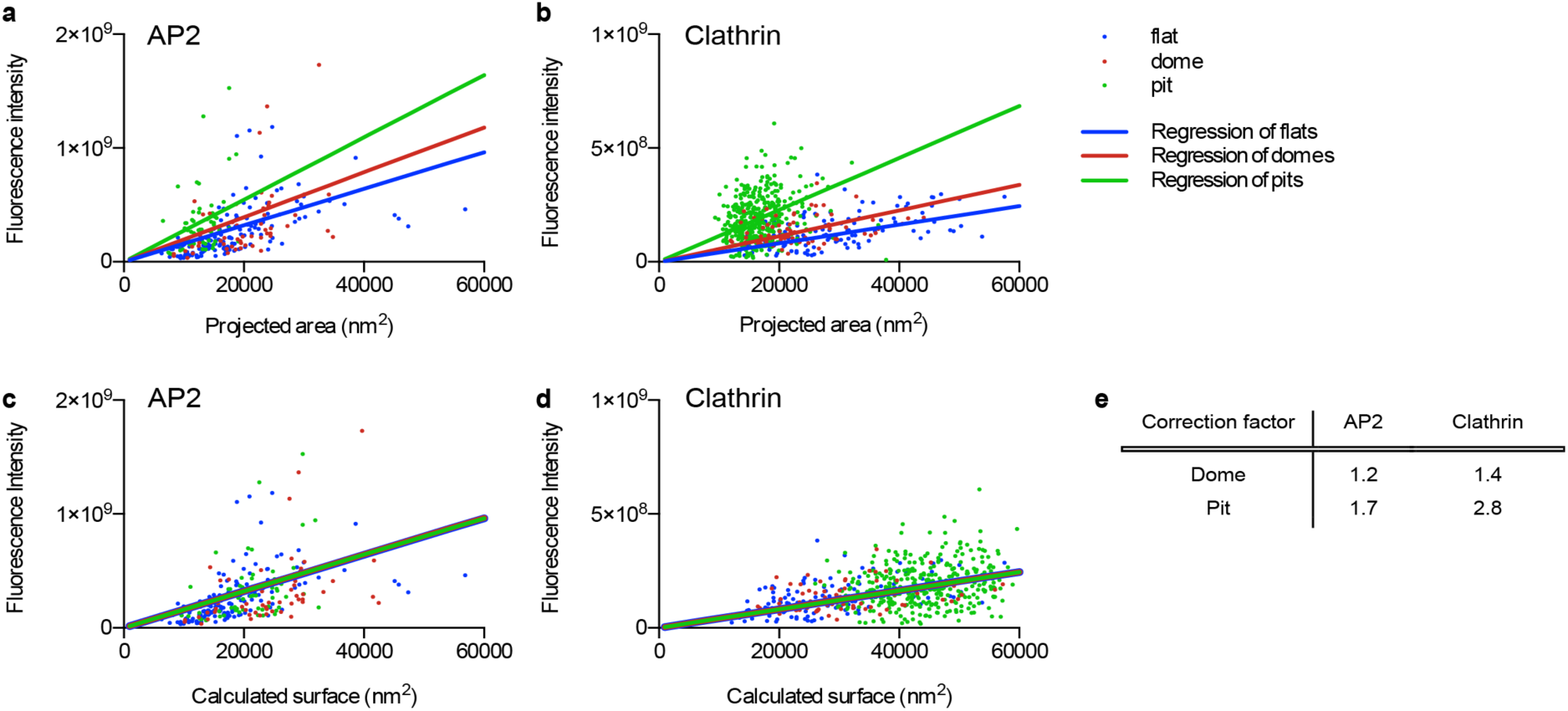
The relative amount of AP2 and clathrin molecules per surface unit of a CCS is curvature dependent. CLEM analysis of CCS labelled with AP2-eGFP (a) or clathrin heavy chain antibody (b). Flat (blue), dome (red), and pit (green). Lines in the corresponding colour show linear regression of the projected area and of the fluorescence intensity. CLEM analysis of CCS corrected according to the regression of flat structures labelled with AP2-eGFP (c) or clathrin heavy chain antibody (d). Projected areas of dome and pit structures of the CLEM analysis were multiplied by a correction factor to fit the linear regression of flat CCS. Lines in the corresponding colour show linear regression of the calculated surface and the fluorescence intensity. (e) Table shows correction factors for dome and pit structures for AP2-eGFP or clathrin heavy chain labelling used in (c) and (d).

As the AP2/clathrin ratio depends on the curvature of CCS (Fig. 4) and as AP2 partitions at different nanoscale zone in relation to the edge of the clathrin lattice during the different stage of CCS formation^22^, we hypothesised that the change in the AP2/clathrin ratio correlates with the stage at which a flat CCS bends to form a CCP. Using cells expressing both AP2 fused to GFP and CLC fused to the fluorescent protein tdtomato, we analysed the intensity profiles of AP2 and CLC during CME. While AP2 profiles often show a distinct plateau phase, the intensity of CLC continues to increase until the end of an endocytic event (Fig. 5a and Supplementary Fig. 1). By normalising the fluorescence intensities of AP2 as well as CLC to the time point when the AP2 signal plateaus, we can calculate the time offset between the time AP2 signal reaches its plateau and the time point CLC reaches its maximal intensity (Fig. 5a-b). Similarly, we defined the intensity offset of clathrin over AP2 (Fig. 5a and c). We found that the time offset was around 10s (Fig. 5b) and that the intensity offset of clathrin over AP2 was around 15% (Fig. 5c). We hypothesised that the time point when AP2 reaches its plateau and therefore the AP2/clathrin ratio changes marks the starting point of bending. We performed another round of CCP assembly modelling, this time using both AP2 and CLC intensity profiles and defining the time point of flat-to-curved transition when AP2 reaches its plateau phase (Fig. 5e and Supplementary Information). At this precise time, the mean clathrin content reached around 70% of its maximal value (Fig. 5d). Using these new parameters, the predicted ratio of flat, dome, and pit structures perfectly matched the EM data (Fig. 5f) and the means of the predicted projected area of both the flat and pit CCS have similar sizes (Fig. 5g-h, box/whiskers and 5i). This AP2/clathrin ratio model better resembles the parameters measured in EM compared to both the constant curvature and constant area models (Fig. 3h and 5i). These findings strongly support a model where flat-to-curved transition initiates at around 70% of its maximal clathrin content and correlates with the concomitant change in the AP2/clathrin ratio.

**Figure 5:**
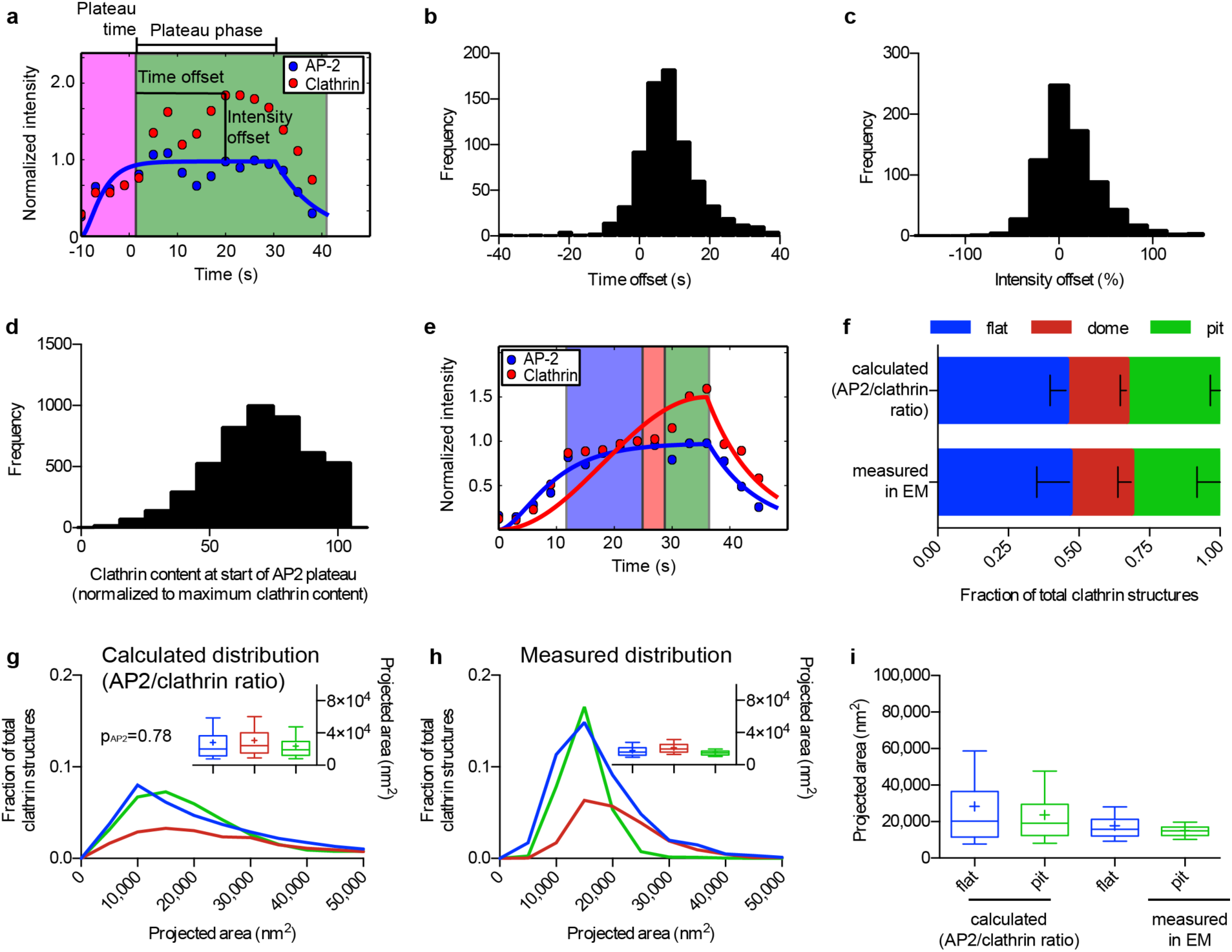
Change in the AP2/clathrin ratio is associated with flat-to-curved transition. (a) Example of an AP2 (blue) and clathrin (red) intensity profile from an individual CME event. The AP2 profile was fitted to equation 1 to find the time when AP2 signal plateaus. For more information see Supplementary Information. The fluorescence intensity of AP2 and clathrin were normalised to the fluorescence intensity of the time when the fitted AP2 profile reaches its plateau. Time offset (difference between the time AP2 plateaus and clathrin reaches its maximum intensity) and intensity offset (excess of maximal clathrin signal over AP2 maximum intensity) are indicated in the profiles. Quantification of the time offset (b) and the intensity offset (c) for 754 tracks of one single cell. (d) Quantification of the clathrin content at the time when AP2 reaches its plateau from 4927 FM tracks. The clathrin signal was normalised to the maximal clathrin signal in each track. (e) Example of an AP2 (blue) and clathrin (red) profile fitted to equation 1 and 2, respectively. These fits where used to calculate the size and curvature distributions of the CCS in (f and g). For more information see Supplementary Information. (f) Comparison of the calculated ratio of flat, dome, and pit structures to the measured ratio in TEM. Results calculated from 4927 FM tracks from 4 different cells; means with SD are shown. (g) Calculated projected area of the CCS using a growth model where the flat-to-curved transition corroborates with the change of clathrin/AP2 ratio (when AP2 signal reaches its plateau phase) for 4927 FM tracks of 4 different cells. P-value of Welch’s t-test to compare the predicted to the measured distribution in panel f. A box/whiskers plot of the projected area is shown in the inset. (h) Measured projected area and curvature distributions of the CCS from TEM data as shown in Fig. 1. A box/whiskers plot of the projected area is shown in the inset. (i) Direct comparison of the projected area distribution of flat and pit structures calculated according to the AP2/clathrin ratio as well as measured in EM, box/whiskers plot.

### High membrane tension inhibits the flat-to-curved transition of CCS

By inducing curvature to the PM, the CCS needs to act against the plasma membrane tension (PMT). Higher PMT has been shown to increase the lifetime of clathrin events at the PM^28,29^ and modelling of the energetic cost of membrane bending suggests that it affects the morphology of the CCS^30,31^. But the effects of increasing PMT on the ultrastructural organisation of CCS have not been investigated in living cells. We monitored the dynamics of clathrin and AP2 during osmotic shock in which the PMT was increased by applying hypotonic medium inducing osmotic swelling of the cells^28^ (Fig. 6a). Following a short latency period, we observed that CCS stalled at the PM. This effect was transient and cells quickly reverted to normal clathrin dynamics (Fig. 6b and d). By performing similar modelling of CCP assembly using AP2 and CLC intensity profiles under osmotic shock, we showed that the CCS display a longer AP2 plateau phase (Fig. 6c and e) and that the time offset was increased compared to mock treated cells (Fig. 6f and 5b). According to our findings that the change in AP2/clathrin ratio coordinates the flat-to-curved transition of CCS, the delayed offset in the AP2/clathrin ratio under higher PMT suggests that the flat-to-curved transition is suppressed and that the coats are flat under this condition.

**Figure 6:**
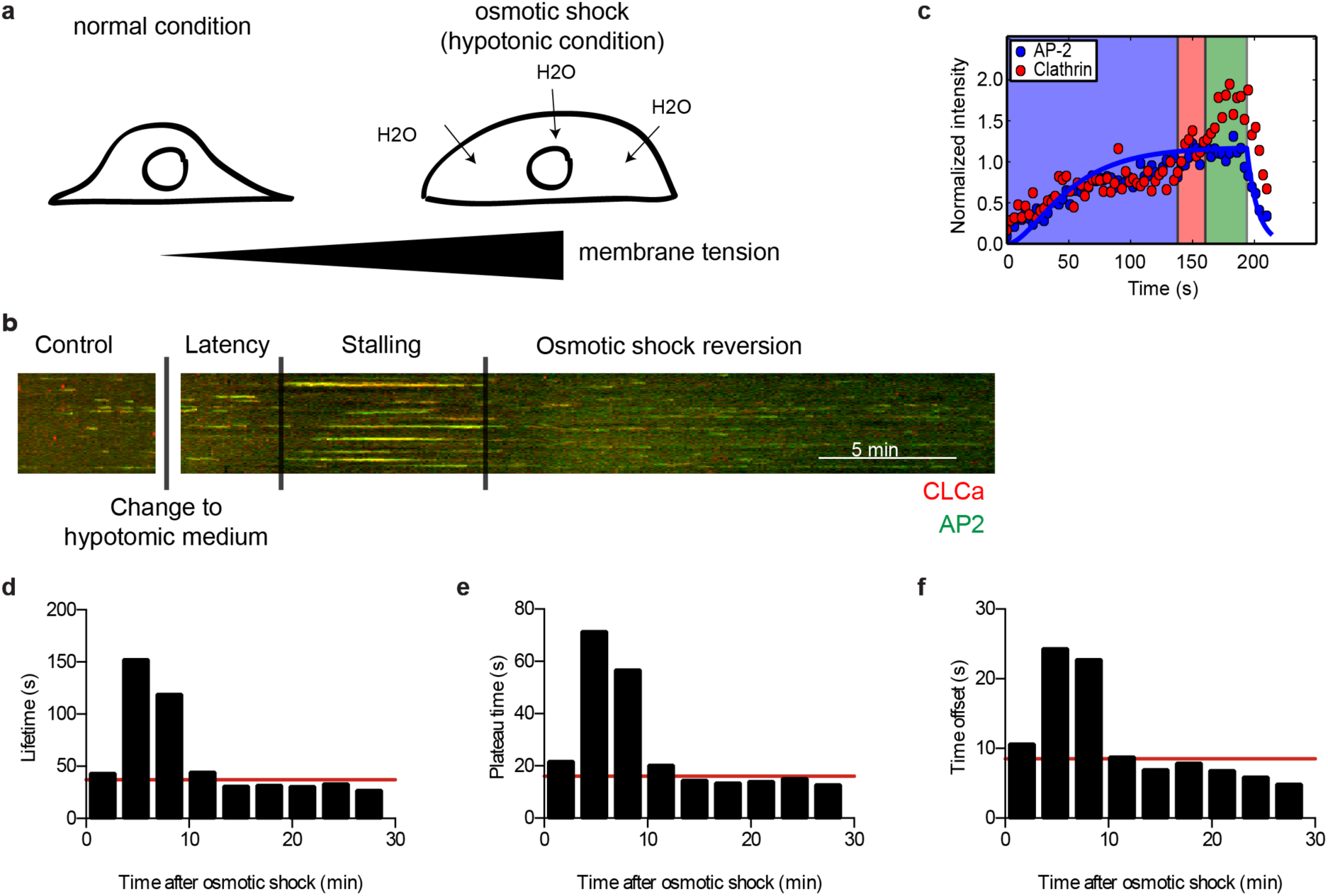
Osmotic shock induces stalling of CCS. (a) Illustration of the effect of osmotic shock on BSC-1 cells. Hypotonic medium was applied to BSC-1 cells, inducing osmotic swelling that results in an increase in PMT. The same BSC-1 expressing fluorescently tagged clathrin light chain and AP2 proteins was followed from 5 minutes prior (internal control) until 30 minutes post hypotonic medium application using spinning disc confocal microscopy. (b) Kymograph of AP2-eGFP (green) and clathrin light chain a-tdtomato (red) expressing BSC-1 cells. The dynamics of CCP was recorded during 5 min prior to osmotic shock until 30 minutes post-osmotic shock. The time after applying the hypotonic medium can be divided into latency, stalling, and osmotic shock reversion time depending on the effect on CME dynamics. Scale bar: 5 minutes. (c) Representative AP2 (blue) and clathrin (red) intensity profile from an individual CME event during the time of stalling fitted to equation 1 to quantify the plateau time. (d) Quantification of the lifetime of CME events during osmotic shock experiments for 1607 tracks of one single cell. CME events were binned in 3 min intervals in respect to the onset of osmotic shock. Red line indicates lifetime of CME prior to osmotic shock. (e) Quantification of the plateau time of AP2 of individual CME events during osmotic shock experiments (as defined in Fig. 5a) for 1607 tracks of one single cell. CME events were binned in 3 min intervals in respect to the onset of osmotic shock. Red line indicates plateau time of CME prior to osmotic shock. (f) Quantification of the time offset between AP2 plateau and clathrin maximum of individual CME events during osmotic shock experiments (as defined in Fig. 5a) for 1607 tracks of one single cell. CME events were binned in 3 min intervals in respect to the onset of osmotic shock. Red line indicates time offset of CME prior to osmotic shock.

By looking at the AP2/clathrin ratio during osmotic shock we predicted that during the stalling phase, 70% of the CCS would be flat (Fig. 7a). To test this notion, we performed EM of metal replica of CCS under osmotic shock (Fig. 7b). We found an accumulation of flat CCS under osmotic shock compared to normal conditions and the frequency was comparable to our predictions from the AP2 and CLC profiles (Fig7c). These flat structures, as well as the dome and pit structures, have the same size distribution as under normal conditions (Fig7d box/whiskers and 7e). The EM data confirms that under higher PMT the flat-to-curved transition of CCS is inhibited. Using mathematical modelling and individual clathrin and AP2 intensity profiles acquired under osmotic shock, we could predict the morphology of the stalled CCS. This validates our model describing that the change in AP2/clathrin ratio represents the precise moment at which the flat-to-curved transition occurs.

**Figure 7:**
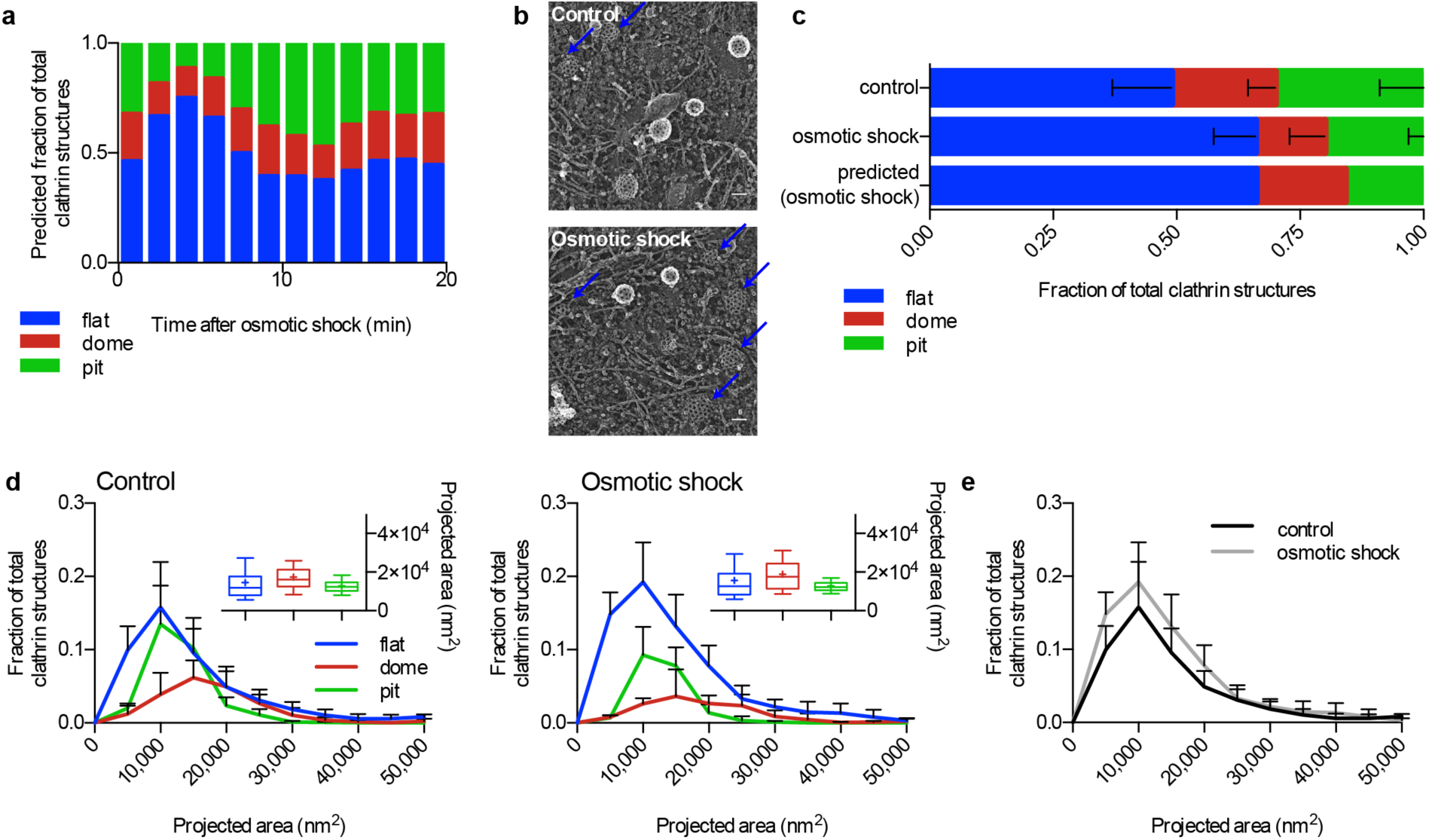
Osmotic shock blocks flat-to-curved transition of CCS. (a) Predicted ratio of flat (blue), dome (red), and pit (green) structures calculated from the binned AP2 and clathrin profiles of CME events (Fig. 6c) during osmotic shock for 1357 tracks. (b) Examples of CCS under normal and osmotic shock conditions. Blue arrows point to flat structures. (c) Comparison of measured and predicted frequency of flat, dome, and pit structures under normal and osmotic shock conditions. (d) Projected area distribution of the different clathrin morphologies under normal and osmotic shock conditions. A box /whiskers plot of the projected area is shown in the inset. Mid-line represents median, cross represents the mean and the whiskers represent the 10 and 90 percentiles. (e) Comparison of projected area distributions of flat CCS under normal and osmotic shock conditions. Results are calculated from four different membranes (number of CCS per membrane: normal conditions 267, 308, 229, and 323; osmotic shock: 395, 99, 351, and 201); means with SD are shown.

## Discussion

The complex coordination of CCS formation during CME has been investigated for decades^32,33^ and the field has been driven by the competition between the constant curvature versus the constant area models^13^. Recent technological advances, in particular in CLEM, favoured the constant area model^19^. In this work, we implemented a multidisciplinary approach to combine information from EM, fluorescence intensity profiles of individual endocytic events and mathematical modelling of CCS growth to infer the ultrastructural rearrangement of CCS during CME.

By modelling the growth behaviour according to the two proposed growth models, we could calculate the expected size and morphology distribution of CCS (Fig. 1 and 3). We could clearly demonstrate that neither of the proposed models explains the ultrastructural organisation and size distribution of CCS present in BSC-1 cells. Instead, our data supports a model in which CCS first grow flat and then the flat-to-curved transition occurs at around 70% clathrin content (Fig. 8). Importantly, we demonstrated that this transition is directly linked to PMT and correlates with a change of the AP2/clathrin ratio within the coat. Increasing PMT results in inhibition of the change in AP2/clathrin ratio and the subsequent stalling of the ultrastructural rearrangement. Our model where a change in AP2/clathrin ratio drives the flat-to-curved transition is consistent with our recent observation that AP2 (and other adaptor/accessory proteins) partitions in different nanoscale area of the clathrin coat and that the concentration of AP2 varies within these zones at various stage of CCS assembly^22^.

**Figure 8:**
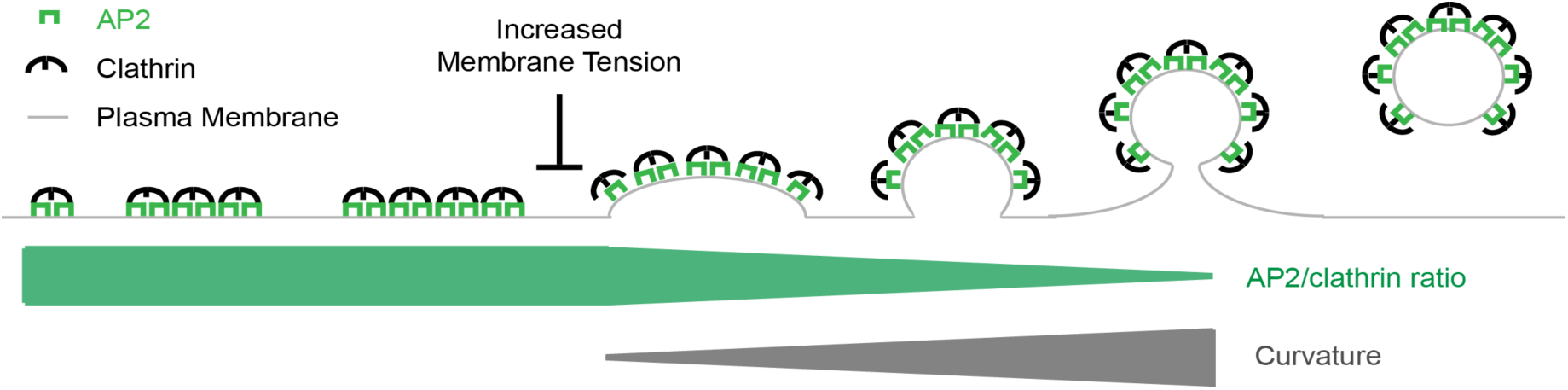
Model of CCP assembly. Schematic representation of the growth model of CCS. CCS initiate as flat clathrin array. They first grow in size in a flat morphology with a constant AP2/clathrin ratio. When they reach around 70% of their full clathrin content, the AP2/clathrin ratio starts to decrease and the CCS start acquiring their curvature. CCPs keep growing by adding additional clathrin molecules until formation and release of CCV into the cytoplasm. The flat-to-curved transition of CCS can be inhibited by increasing PMT resulting in an accumulation of flat structures. We propose that flat-to-curved transition is concomitant with bypassing the energy barrier necessary to curve the PM and that this critical step in CME is coordinated by the uncoupling of clathrin and AP2 characterised by their abrupt ratio decrease.

Our growth behaviour of CCP assembly, determined in this work in BSC-1 cells, may also apply for other cell types. To test whether our growth model could explain the ultrastructural distribution of CCS described in previous studies, we extended our mathematical model and calculated the predicted contact angle between the clathrin cage and the PM, the coated surface area and the radius of tip curvature of the CCS (Supplementary Fig. 3). By comparing the model to the Avinoam et al. data^19^ set describing the distribution and ultrastructural organization of CCS in SK-MEL-2 cells, we could demonstrate that our growth model could also explain their observed distribution.

It was proposed earlier that the physical properties of the PM are influencing the morphology of CCS because PMT energetically acts against curvature acquisition of the clathrin array^28,30,31^. We demonstrated the effect of increased PMT on CCS assembly in cells. Under high PMT, the accumulation of stalled flat CCS at a stage prior of the change in AP2/clathrin ratio reveals an important step of CCP formation to overcome the PMT to acquire curvature. It is tempting to speculate that this change of AP2/clathrin ratio is a key event mandatory for curvature acquisition. The demonstrated interplay between AP2 and PMT shows that both biochemical and physical factors regulate CME. Surprisingly, the increase in PMT only changes the ratio between the different morphologies of CCS in favour of flat structures but does not affect their size. One could assume that a higher energy barrier of the PM might be counteracted by the clathrin system by the formation of larger flat CCS, which could accumulate more energy for the bending process by molecular crowding of clathrin itself as well as BAR proteins incorporated in the coat^30,34–36^. Since under high PMT the flat CCS still have the same size as under normal PMT we suggest that there is an internal limitation of the coat size that might be regulated by certain components of the coat. Proteins that regulate the size of CCV have been reported^37,38^ and it might be possible that the size of a flat clathrin lattice is controlled in a similar way. The fact that other cell types show much larger flat CCS under normal conditions, commonly referred to as clathrin-coated plaques^10,15^, illustrates that the clathrin machinery is capable of forming large flat structures under certain conditions. But the factors necessary for clathrin-coated plaque formation have not been described so far. These clathrin-coated plaques might contribute to CME. Live-cell FM^39,40^ and EM^41^ of such structures support budding of CCV from such plaques most probably from the edge, again illustrating the ability of a flat CCS to rearrange into a CCP further supporting that our observed flat-to-curved transition is indeed possible.

We could show that during coat assembly the AP2/clathrin ratio changes. This finding is in agreement with other FM^26,27,29^ and CLEM^22^ studies. Mathematical modelling demonstrated the correlation between the change in the AP2/clathrin ratio and the time of coat bending. Several other adaptor and accessory proteins have been proposed to influence the ultrastructure of the CCS. Depletion of FCHO 1 and 2 has been reported to alter the ordered hexagonal organisation of the flat clathrin lattices^42^. Other proteins like CALM and NECAP have been proposed to regulate the final size off a CCV^37,38^.

Our findings provide a unifying view on the process of CCP assembly where we demonstrate that CCPs initially grow as flat lattices and that change in clathrin/adaptor ratio correlates with the onset of coat curvature acquisition prior to the completion of coat polymerization. We propose that the proportion of different coat proteins could ultimately define the morphology of a clathrin structure and temporal changes of this proportion might initiate bending of the coat, allowing for dynamical regulation by the cell.

## Author contributions

DB and KAS performed CLEM experiments. DB performed FM experiments. FF performed modelling. DB and FF designed experiments and modelling, analysed data and wrote manuscript. KAS performed EM experiments. SK and H-GK performed STED. J-PB, WJG and KR tracked FM data. JWT designed experiments and oversaw EM and CLEM imaging. USS designed modelling, interpreted data and wrote manuscript. SB designed experiments, interpreted data and wrote manuscript.

### Acknowledgments

This work was supported by a research grant from Chica and Heinz Schaller Foundation and Deutsche Forschungsgemeinschaft (DFG) in SFB1129 (Project 14) to SB. DB was supported by a fellowship of the Hartmut Hoffmann-Berling International Graduate School of Molecular & Cellular Biology (HBIGS) at the Heidelberg University and by a travel grant from Boehringer Ingelheim Fonds. FF was supported by a fellowship of the Heidelberg Graduate School of Fundamental Physics (HGSFP) at the Heidelberg University. USS and SB are members of the cluster of excellence CellNetworks. J. W. Taraska was supported by the Intramural Research Program of the National Heart Lung and Blood Institute, National Institutes of Health, USA. We would like to thank Ulrike Engel and the Nikon Imaging Center (Heidelberg University) for support with TIRF microscopy, Vibor Laketa from the Department of Infectious Diseases, Virology (University Hospital Heidelberg) for support with spinning disc microscopy, and the US National Heart Lung and Blood Institute (NHLBI) Electron Microscopy Core and Light Microscopy Core facilities for use of equipment.

## Methods

### Cell lines and cell culture

BSC-1 cells were obtained from ATCC. BSC-1 cells stably expressing AP2-eGFP were created by transfecting BSC-1 cells with a plasmid expressing the sigma2 subunit of AP2 fused to eGFP^20^. Following selection with G418 (750 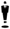 g.mL^−1^) (Gibco), AP2-eGFP expressing BSC-1 cells were grown in DMEM (Gibco) supplemented with 10% fetal bovine serum, penicillin and streptomycin (Gibco) at 37 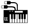 C and 5% CO_2_. For passaging, cells were rinsed with PBS and incubated for 3-5 minutes with 0.05% Trypsin/EDTA (Gibco) at 37 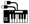 C and 5% CO_2_. After detaching, the cells were resuspended in complete medium. Passaging was done every 2-3 days in a ratio of 1:5-1:10.

### Transfection

Transfection of cells was done using Lipofectamine 2000 (Invitrogen). Cells were plated in 6-well plates one day before transfection. The next day, cells were transfected at 70-80% confluence. 2μg DNA and 8μl Lipofectamine 2000 were separately mixed with 100μl OptiMEM (Gibco). The two solutions were mixed together. After incubation for 20 minutes at room temperature, the transfection mix was added drop-wise onto the cells. For generation of stable cell lines, the growth medium was exchange for fresh growth medium after 8 hours. The cells were put under selection two days after transfection.

For live-cell imaging of BSC-1 AP2-eGFP transiently expressing CLCa-tdtomato were seeded 8 hours after transfection.

### Antibodies and Plasmids

Anti-clathrin heavy chain antibody (X22, ab2731, 1:500 for immunofluorescence) was purchased from Abcam. Anti-clathrin light chain antibody (Con.1, C1985, 1:500 for immunofluorescence) was purchased from Sigma-Aldrich. Secondary Alexa

Fluor 647 goat anti-mouse (1:1,000 for immunofluorescence) and goat anti-mouse Atto594 (1:200 for immunofluorescence) were purchased from molecular probes. Wheat Germ Agglutinin AlexaFluor 647 conjugate (W32466, 1:200 for immunofluorescence) was purchased from molecular probes.

Mammalian expression vectors containing rat CLCa N-terminally fused to tdtomato^43^ and the rat AP2 subunit sigma2 C-terminally fused to eGFP^20^ were used for stable and transient expression of fluorescently tagged proteins.

### Live-cell microscopy

Glass coverslips (TH. Geyer, 25mm diameter, No. 1.5H) were coated with poly-D-lysine solution (Sigma-Aldrich, #P6407) at concentration 0.1mg ml^−1^ for 5 minutes at room temperature and washed three times with PBS. Cells were seeded on poly-D-lysine-coated coverslips and live-cell microscopy was performed 12-16 hours after seeding. Live-cell imaging of AP2-eGFP was performed with an inverted spinning-disk confocal microscope (PerkinElmer), with a 60x (1.42 numerical aperture, Apo TIRF, Nikon) or 100x (1.4 numerical aperture, Plan Apo VC, Nikon) oil immersion objective and a CMOS camera (Hamamatsu Ocra Flash 4). An environment control chamber was attached to the microscope to keep 37 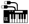 C and 5%CO_2_. 10 minutes-long movies of representative cells were taken with one frame every 3 seconds. Live-cell imaging of AP2-eGFP together with CLCa-tdtomato was performed with an inverted Ti microscope (Nikon) with objective TIRF illumination, with a 60x (1.49 numerical aperture, Apo TIRF, Nikon) oil immersion objective and EMCCD camera (Andor iXon Ultra DU-897U). An on-stage incubation chamber was used to keep 37 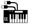 C and 5%CO_2_. 10 minutes-long movies of representative cells were taken with one frame every 3 seconds.

### Osmotic shock experiments

For live-cell microscopy of cells under osmotic shock, one cell was imaged for 5 minutes under normal conditions. Afterwards the medium was change to hypotonic medium (1:1 ratio of medium to water with 10% FBS) and the same cell was imaged for an additional 30 minutes. For TEM of cells under osmotic shock, cells were put into hypotonic medium and unroofed after 10 minutes of osmotic shock. The samples were then prepared for TEM.

### Unroofing

Cells were seeded on poly-D-lysine-coated coverslips (25mm). Unroofing was performed 16 hours after seeding. Cells were washed three times with stabilisation buffer (30mM HEPES buffer, brought to pH 7.4 with KOH, 70mM KCl, 5mM MgCl_2_). Unroofing was performed in 2% paraformaldehyde in stabilisation buffer using two short sonication pulses. Sample was immediately put into fresh 2% paraformaldehyde solution and fixed for 20 minutes at room temperature.

### Immunofluorescence

For immunofluorescence, intact cells growing on poly-D-lysine coated 12mm coverslips (#1.5, Thermo Scientific) or unroofed PMs were fixed with 2-4% paraformaldehyde for 20 minutes at room temperature. Intact cell were permeabilised with 0.5% Triton X in PBS for 15 minutes. After blocking with PBS 1% BSA for 1 hour at room temperature, samples were incubated with primary antibody diluted in PBS with 1% BSA for 1 hour at room temperature. After five washes with PBS, samples were incubated with the secondary antibody. Wheat germ agglutinin diluted in 1% BSA in PBS was incubated on cells for 30 minutes at room temperature. After five washes with PBS, samples for STED were mounted using Mowiol and samples for correlative light and electron microscopy were fixed by incubation with 2% paraformaldehyde for 20 minutes at room temperature and washed three times with PBS.

### Imaging of unroofed PMs for CLEM

Widefield fluorescent images of unroofed PMs were taken with a Nikon N-STORM microscope with a 100x oil immersion objective and an EMCCD camera (Andor Ixon Ultra DU-897). To cover an area of 1mm^2^ a montage of 15x15 images with an overlap of 15% for stitching was taken. The imaged area was marked with a circle (4mm in diameter) around the centre of the imaged area using an objective diamond scriber. The immersion oil was carefully removed from the bottom of the glass coverslip and the sample was prepared for EM.

### TEM of metal replica

Coverslips with unroofed membranes were fixed with 2% glutaraldehyde in PBS overnight. Samples were incubated with 0.1% tannic acid for 20 minutes at room temperature. After four washes with water, the samples were incubated with 0.1% uranyl acetate for 20 minutes at room temperature. After two washes with water, samples were dehydrated with a series of ethanol solutions (15%-100%). Samples were placed in each ethanol solution for 5 minutes. After replacing the 100% ethanol solution, the samples where dried in a critical point dryer. The samples were then put under vacuum until they were coated. The samples were coated with JFDV JOEL Freeze Fracture Equipment with a first layer of platinum with an angle of 17 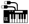 while rotating and with a second layer of carbon with an angle of 90° while rotating. For better orientation, the marked area of the coated samples was imaged with a phase contrast microscope. The samples were then cut to fit on the EM grids (TED PELLA, 75 Mesh Copper, Support Films Formvar/Carbon). 5% hydrofluoric acid was used to remove glass from the metal replica. The floating metal replica was extensively washed with water and the carefully placed on a glow discharged EM grid. Samples were dried on filter paper and again imaged with a phase contrast microscope. TEM imaging was performed using a JEOL 1400 equipped with SerialEM freeware for montaging. Montages of large membrane sheets at 12,000 magnification (1.82nm per pixel) with 10% overlap were imaged.

### Transformation of images for CLEM

The fluorescence microscopy image and the electron microscopy montage of the same membrane sheet were first manually and roughly overlaid using Photoshop. MATLAB was used to transform the fluorescence image according to the electron microscopy montage using three manually identified CCS. For the transformation the centre of the clathrin structure in the electron microscope montage and the centre of the fluorescence signal determined by a Gaussian fit were used as landmarks.

### Tracking

For tracking CME events, we used ilastik (http://ilastik.org). First the images were segmented using the pixel classification and object classification workflow. For tracking, the automatic tracking workflow was used. The maximal distance was put to 5 to avoid merging of close tracks. Automatic tracking of CCS using AP2-eGFP expressing BSC-1 cells was performed using a probabilistic particle tracking method^44^. The method combines Kalman filters with particle filters and probabilistic data association with elliptical sampling (PDAE). For particle detection, a Laplacian-of-Gaussian filter and connected-component labelling was used. Based on the computed trajectories, the signal intensity of each tracked object (normalized to the background intensity) was determined and intensity statistics over all trajectories were computed. Also, the object size was determined for each time. In addition, the lifetime of CCS was quantified and classified into different ranges.

### STED

STED nanoscopy was carried out using the Two-color-STED system (Abberior Instruments GmbH, Göttingen). Image acquisition was performed using a 100x Olympus UPlanSApo (NA 1.4) oil immersion objective and 70 % nominal STED laser power (λ= 775 nm, max. power = 1.2 W). Line accumulation was set to 4 and a pixel size of 15 nm was used. Deconvolution of acquired images was done using imspector software (Abberior Instruments GmbH, Göttingen). Richardson-Lucy deconvolution with a regularisation parameter of 0.001 was used and stopped after 30 iterations.

## Supplementary Information

### Constant area model

As the fluorescence intensity of labelled clathrin triskelia is proportional to the number of incorporated clathrin triskelia or equivalently to the size of clathrin covered membrane area, we model the assembly of CCV as surface growth.

Our mathematical description of the constant area model considers finite growth of a flat circular patch with radius r. The flat patch grows at its edge (L) with rate *K*_on_ but decays over the bulk with rate *K*_off_, which can be expressed by 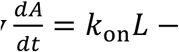. In (Fig. 3a) a sketch of the growth schematics is shown. The growth equation can be simplified by plugging in the surface of a circular patch *A* = *πr*^*2*^., it follows 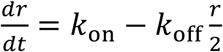 with steady state radius *r*_ss_ = 2*K*_on_/*K*_off_. By integrating this equation, we find the patch area as a function of time

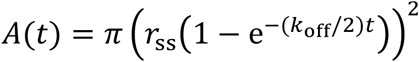

In the observed fluorescence intensity tracks the intensity decreases after some time until the intensity vanishes completely. Biologically, this indicates that the clathrin coated vesicle pinches off the cell membrane and therefore, moves out of the focus of the microscope. We model this by assuming an exponential decay of the area with time constant *τ*, starting at time *t*_decrease_

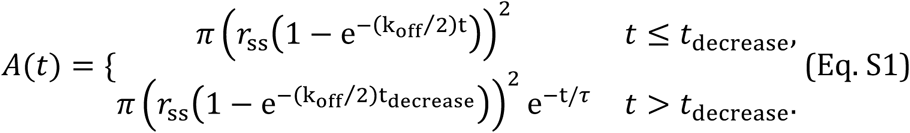

As the steady state is only reached approximately we define an area plateau at 95% of the steady state area and the corresponding time *t*_plateau_. The constant area model assumes that a flat patch transforms into a spherical pit as soon as the patch reaches the area plateau. Here, we neglect the exact details of the transformation process but classify the CCS as flat, dome (less than a half sphere) or pit according to time. Before reaching the plateau, the area is considered flat. After reaching the plateau we classify CCS in the first 40% of the remaining time until *t*_decrease_ to be domes and in the last 60% of time to be pits. The exact choice is motivated by the occurrences of domes and pits in EM which is roughly 2:3.

### Data fits

To test whether the constant area model correctly describes the shape and size of clathrin coated vesicles we fitted Eq. 1 to 4927 FM tracks of 4 different cells (Fig. 3b) and calculated from the fitted area surface growth curves histograms which we could compare to EM histograms. Therefore, we related the intensity of an FM track to its corresponding area. Furthermore, the FM dataset was filtered before the fitting. The exact details of our procedure are described in the following.

### Relate fluorescence intensity and surface area by means of CLEM

To relate the fluorescence intensity of a clathrin FM track to the corresponding clathrin covered membrane area we use our clathrin CLEM data, relating the projected surface size of CCS to their fluorescence intensity. We analyse flat CCS for which the projected area directly corresponds to their surface. By fitting a line through the origin to the clathrin CLEM data we get the slope *β* providing us with a linear relation between size and intensity. As the local intensity background is removed from the CLEM data we find

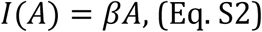

where I is the intensity of the FM data, A is the area of the clathrin structure and *β* is the proportionality constant. Fig. 4b shows the intensity of flat structures as a function of their projected area (blue). We find *β* = 4085*nm*^*2*^.

### Relate different FM datasets

To calculate a size histogram from fluorescence intensity tracks we analyse live cell FM data, that have a different intensity level than the fluorescence intensity of the CLEM data. Therefore, we need to relate these two different data sets. As the CLEM intensity I and the live cell FM intensity I' are both proportional to the number of labelled clathrin triskelia, both intensities, which are background corrected, can be related only by some factor *α*

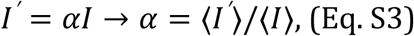

which can be calculated by dividing the means (indicated by 〈〉) of both data sets.

### Relate different FM datasets

We next restricted the live cell FM data set to ensure that the calculated size histogram is comparable to the EM histogram. In EM we only detect structures that reach a threshold size. However, in the live cell FM data set we register only detectable intensities, which exceed the local background signal. Therefore, we relate the minimal detectable size in CLEM (*A*_*T*_) to a threshold intensity (*I*_*T*_), which we relate to a threshold intensity (*I*^*′*^) that we use on the live cell FM tracks. As *I*^*′*^ = *αI* we find

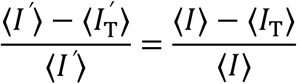

As 〈*I*^*′*^〉 is a function of 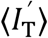 we can find the root of

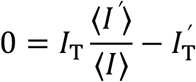

by iteratively increasing 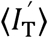.

In EM we find that the minimal sized clathrin structure has an area of *A*_*T*_ = 5644*nm*^2^., corresponding to a calculated intensity of *I*_*T*_ = 2.305. 10^7^arb. unit. We find the mean intensity in the clathrin CLEM data of all structures 〈*I*〉 = 1.766. 10^8^arb. unit. To measure the mean intensity in the live cell FM data set we sample from each FM track that lasts a least 24 seconcds a number of intensities proportional to its lifetime. From these sampled intensities, which we restrict to be larger than the threshold value *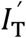*, we then compute the mean intensity. We obtain *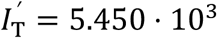*, 〈*I*^*′*^〉 = 4.248. 10^4^arb. unit. By combining the Eq. S2 and Eq. S3 we finally arrive at

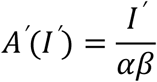

according to which we compute the surface of a clathrin structure from its intensity in the live cell FM. Furthermore, we filter all FM tracks showing a mean intensity which is smaller than 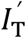, as those tracks would not be observable in EM.

### Data filtering

To ensure that all structures correspond to real objects we assume for a productive object a minimal lifetime of 24 seconds. We filter our data set for FM tracks with multiple structures (defined as a FM track that shows at least two clear intensity maxima) to allow for direct fitting of single tracks. Furthermore, we filter tracks that start already with a medium intensity level *I*^*′*^(*t* = 0) > 0.5〈*I*^*′*^〉.

### Parameter choice and data fitting

The parameters for the fit are restricted by assuming that growth curves should at least reach 90% and at most 120% of the maximal area value. Additionally, we assume that vesicle pinch off (corresponding to a decrease in the intensity to 10%) takes between 10 seconds to 20 seconds. Additionally, we require the fit to reach 99% of the steady state area before the area decrease happens and at least 10% of that time until the 99% area level are reached. We implement the Python module ‘lmfit’ for fitting the area tracks where we use the method ‘nelder’ of the minimiser function. In this way we obtain for each track four parameters (*r*_ss_, *K*_off_, *τ*, *t*_decrease_) that characterise the growth curve.

### Calculation of the size histogram

From the growth curves, we calculated a histogram (Fig. 3c) to compare the constant area model to the measured projected size EM histogram (Fig. 3h). We proceeded as follows:

- From each of the fitted growth curves we uniformly drew a number of time points, proportional to the time until the structure pinches off the cell membrane, that is given by *t*_decrease_ (Fig. 3a and b). We used 4927 FM tracks of 4 different cells giving us 4 million time points, which we used to calculate the size histogram.
- We classified the chosen times and corresponding areas into three categories. If *t* < *t*_plateau_ the structure is flat (blue region in Fig. 3b). We assume that the transformation process from flat to dome takes 40% of the plateau time (which corresponds roughly to the ratio between domes and pits in EM) whereas from dome to pit it takes 60%. This is supported by the notion that the clathrin array reorganisation becomes increasingly complicated. Therefore, if *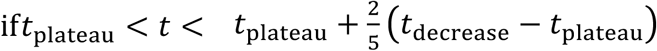* the structure is a dome (red region in Fig. 3B) and otherwise it is classified as a pit (green region in Fig. 3b).
- We computed the projected area by assuming that the transformation within the dome and pit phase is a linear function of time. Therefore, we divided the area by 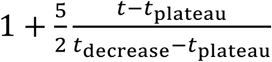 for domes and 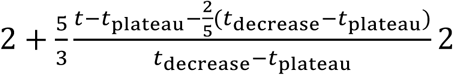 2 for pits. This factor equals 1 for a flat patch, 2 for a completed dome (half sphere) and 4 for a completed pit (full sphere).
- We excluded times for which the corresponding area is below the area threshold or where the area already decreases (white region in Fig. 3b).
- From the calculated projected areas, we then determined the size histogram (Fig. 3c), which we compare to the EM data (Fig. 3h).

### Curvature acquisition during growth: updated model

In the updated growth model we assume that CCS first grow flat, start to invaginate as they reach 70 % of their final size (which we determine by taking the inverse of the intensity ratio of pit and flat structures in clathrin CLEM, which is 1.44) and finally grow as a spherical cap until a full pit has formed. As before we model the assembly of clathrin coated vesicles as surface growth.

Our mathematical description of the “curvature acquisition during growth model “ considers a spherical cap with radius *R* that grows at its edge with rate *K*_on_ which can be expresses by 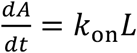. In (Fig. 3d) a sketch of the growth schematics is shown. The growth equation can be simplified by plugging in the surface of a spherical cap *A* = 2*πR*. ^2^ (1– cos(*φ*)), such that the change in the angle 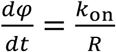 is constant. In the limit of a small growth angle *φ* = *K*_on_*t*/*R* < 2 we recover the growth equation of a flat patch *A*(*t*) = *π* (*K*_on_*t*)^2^. For an almost complete pit the growth equation also holds perfectly. However, the mathematical description of the growth model approximates the flat patch by a spherical cap for intermediate flat patch sizes. In this case, the error is negligible as the model interpolates between the correctly addressed limiting cases and is not used to assign the shape of the CCS.

To define the ultrastructural organization of the clathrin lattice, we neglect, as before, the exact details of the transformation process but classify the CCS as flat, dome (less than a half sphere) or pit according to time relative to the time when reaching the maximal area. Before reaching 70% of the maximal area the coat is flat. After reaching 70% of the maximal area we classify CCS in the first 40% of the remaining time to be domes and pits otherwise. The exact choice is motivated by the occurrences of domes and pits in EM which is roughly 2:3.

As before we model that vesicles pinch off the membrane by an exponential decay of the area with time constant *τ*, starting at time *t*_decrease_ = *πR*/*K*_on_. The full growth equation for the area as a function of time *A*(*t*) reads

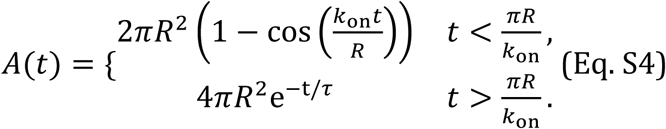

### Data fits, parameter choice and data fitting

To test whether the “curvature acquisition during growth” model correctly describes the shape and size of clathrin coated vesicles we fitted Eq. S4 to 4927 FM tracks of 4 different cells (Fig. 3e) and calculated from the fitted area surface growth curves histograms which we could compare to EM histograms. Therefore, we related the intensity of an FM track to its corresponding area. Furthermore, the FM dataset was filtered before the fitting. The exact details of our procedure are the same as before.

The parameters for the fit are restricted by assuming that growth curve should at least reach 90% and maximal 120% of the maximal area value and that the vesicle pinching off (corresponding to a decrease in the intensity to 10%) takes between 10 seconds to 20 seconds. We implement the Python module ‘lmfit’ for fitting the area tracks where we use the method ‘nelder’ of the minimiser function. In this way we obtain for each track 4 parameters (*R*, *K*_”$$_, *τ*,) that characterise the growth curve.

### Calculation of the size histogram

From the growth curves we calculated a histogram (Fig. 3f) to compare “the curvature acquisition during growth model” to the measured projected size EM histogram (Fig. 3h). We proceeded in principle as before and only mention changes:

- We classified the chosen times (around 4 million time points) and corresponding areas into three categories. If *A*(*t*) < 0.7*A*_max_ the structure is flat (blue region in Fig. 3e) and we call this time *t*_transformation_. We assume that the transformation process from flat to dome takes the first 40% of the remaining time until the maximal area is reached (which corresponds roughly to the ratio between domes and pits in EM) whereas from dome to pit it takes the rest of the time.
- We computed the projected area by assuming that the transformation within the dome and pit phase is a linear function of time. Therefore, we divided the area by 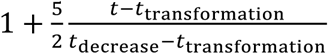 for domes and 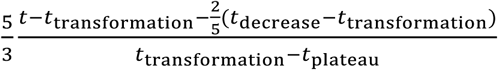. This factor equals 1 for a flat patch, 2 for a completed dome (half sphere) and 4 for a completed pit (full sphere).

### Curvature acquisition during growth: Flat-to-curved transition corroborates with the change of clathrin/AP2 ratio

Here, the only thing that changes compared to the updated model is that the transition time *t*_transformation_ is now given by the time when the AP2 intensity plateaus, which we calculated by fitting eq. 1 to the AP2 FM intensity tracks.

### Live cell FM analysis resampling

To determine the ratio of clathrin and AP2 during the process of CME, we performed TIRF microscopy of BSC-1 AP2-eGFP cells transiently expressing CLCa-tdtomato. We analysed the excess of clathrin in comparison with AP2 during the formation of CCVs. Therefore, we calculated the ratio of the maximum clathrin intensity divided by the intensity of clathrin at the time when AP2 shows an intensity plateau (95% intensity level) and subtract this ratio from the ratio, which we get for AP2.

In detail: we set a threshold for the AP2 intensity underneath fluorescence intensity tracks are excluded (as before). We calculate the plateau time, defined as the time when 95% of the AP2 plateau intensity is reached by fitting the constant area model to the AP2 FM data. Additionally, we normalise all intensity values to the intensity value when 95% of the AP2 intensity (clathrin intensity) is reached. For each track (in total 754 tracks of one single cell) we calculate the difference of the time when AP2 plateaus and clathrin reach the maximum intensity and determine the corresponding histogram (cf. Fig 5b). Furthermore, we calculate the intensity offset given by the normalised maximum clathrin intensity divided by the normalised maximum AP2 intensity (cf. Fig. 5c).

### Calculation of the ratio histogram during the osmotic shock

To determine the ratio histogram of flat, dome and pit CCS during the osmotic shock (Fig. 7a and 7c) we first defined a transition time *t*_transformation_, when flat CCS start to invaginate, where the normalised clathrin intensity exceeds the normalised AP2 intensity by 5% (blue region in Fig. 6c). We assumed that the transformation process from flat to dome takes the first 40% of the remaining time until the vesicle pinches of the PM, given by *t*_decrease_ (red region in Fig. 6c) whereas from dome to pit it takes the remaining 60% of time (green region in Fig. 6c). Next, we started 50s after the osmotic shock and determined the morphology of all tracks present at that time depending on the description above. We repeated this procedure in steps of 5 seconds and average the number of structures over time intervals of 100 seconds. In total we used 1356 FM tracks of one cell. The found ratios of flat, dome and pit CCS were then plotted as a function of the time after the osmotic shock (Fig. 7a).

To test the consistency of this approach we used it to calculate the ratios for flat, dome and pit structures on the data without osmotic shock consisting of 4927 FM tracks of 4 different cells. Averaging over all tracks and considering only tracks with lifetimes shorter than 90s and with AP2/clathrin discrepancy we obtained 47.8% flat CCS, 18.0% dome CCS and 34.2% pit CCS which is close to the determined ratios in (Fig. 5g).

### Quantification of agreement between measured and predicted size histograms

To determine the level of agreement between measured (Fig. 3h) and predicted size histograms (Fig. 3c, Fig. 3f and Fig. 5e) we calculated chi-squared 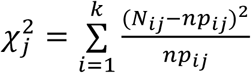 where we sum over all bins *K* the occurrences *N*_*i*_ of measured CCS and compared it with the number of expected occurrences *np*_*i*_, where *p*_*i*_ is the predicted normalised frequency per bin, which we deduced from our models (CAM=constant are model, UM=updated model and AP2=transition flat/dome determined by the time when the AP2 intensity plateaus) and *n* = ∑*N*_*i*_. We repeated this for all CCS *j* = {*flat*, *dome*, *pit*} such that we found three values for 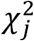. for each model and cell. We averaged these values for all CCS and four cells and found: 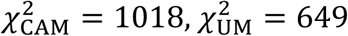 and 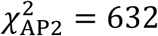 which shows that the model plotted in Fig. 5e describes the data best.

Additionally, we performed a Welch’s t-test to calculate p-values for the null hypothesis that the measured and predicted size distributions of a CCS have identical mean values. We averaged over all CCS and four cells and found *p*_CAM_ = 0.60, *p*_UM_= 0.75 and *p*_CAM_ = 0.78.

---

**Supplementary Figure 1:**
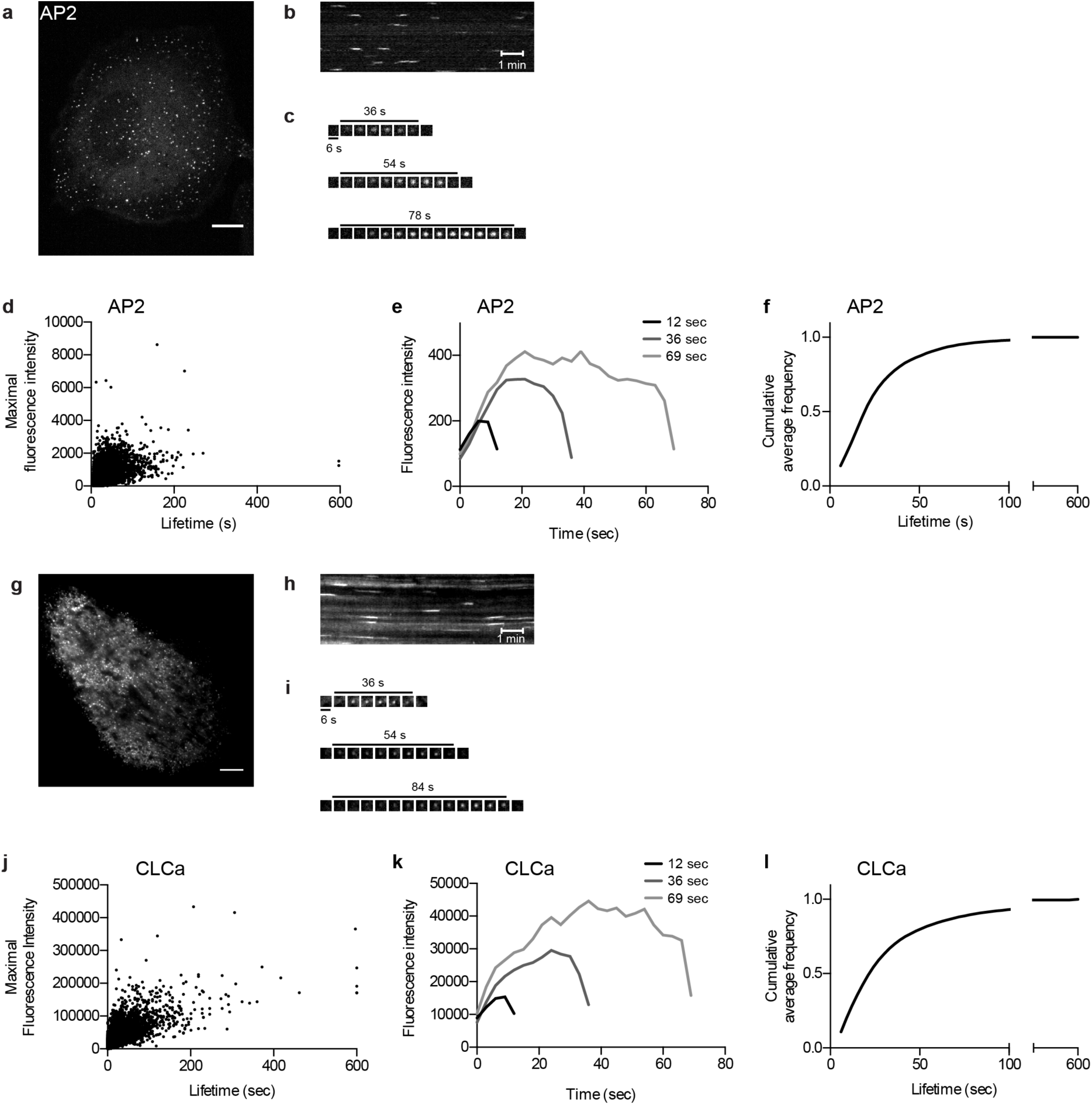
Characterisation of clathrin dynamics in BSC-1 cells. Spinning disc live-cell microscopy of BSC-1 AP2-eGFP cells. (a) Representative image of a BSC-1 AP2-eGFP cell. Scale bar: 10μm. (b) Kymograph of a 10 minute long movie following AP2-eGFP signal. Scale bar: 1min (c) Representative individual CME events. (d) Dot plot showing the lifetime and the maximal fluorescence intensity of AP2-eGFP of all CME events from one cell in a 10 minute long movie. Total number of tracked CME events: 14,694. (e) Average fluorescence intensity profiles (AP2-eGFP) of CME events with lifetime of 12, 36, and 69 seconds. Data are acquired from one cell during a 10 minute long movie. Number of tracks used for calculating the average intensity profiles: 1,525 for 12sec-lifetime; 604 for 36sec-lifetime; 230 for 69sec-lifetime. (f) Cumulative average frequency of CME lifetime. The average was calculated from five different cells. Number of total tracks per cell: 14,037; 16,386; 14,694; 16,797; 19,631. (g) TIRF live-cell microscopy of BSC-1 cells expressing clathrin light chain a (CLCa)-tdtomato. Representative image of a BSC-1 cell expressing CLCa-tdtomato. Scale bar: 10μm. (h) Kymograph of a 10 minute long movie following CLCa-tdtomato. Scale bar: 1min (i) Representative individual CME events. (j) Dot plot showing the lifetime and the maximal fluorescence intensity of CLCa-tdtomato of all CME events from one cell in a 10 minute long movie. Total number of tracked CME events: 4,763. (k) Average fluorescence intensity profiles (CLCa-tdtomato) of CME events with lifetime of 12, 36, and 69 seconds. Data are acquired from one cell during a 10 minute long movie. Number of tracks used for calculating the average intensity profiles: 175 for 12sec-lifetime; 75 for 36sec-lifetime; 21 for 69sec-lifetime. (l) Cumulative average frequency of CME lifetime. The average was calculated from four different cells. Number of total tracks per cell: 2,429; 4,763; 5,443; 16,797; 8,393.

**Supplementary Figure 2:**
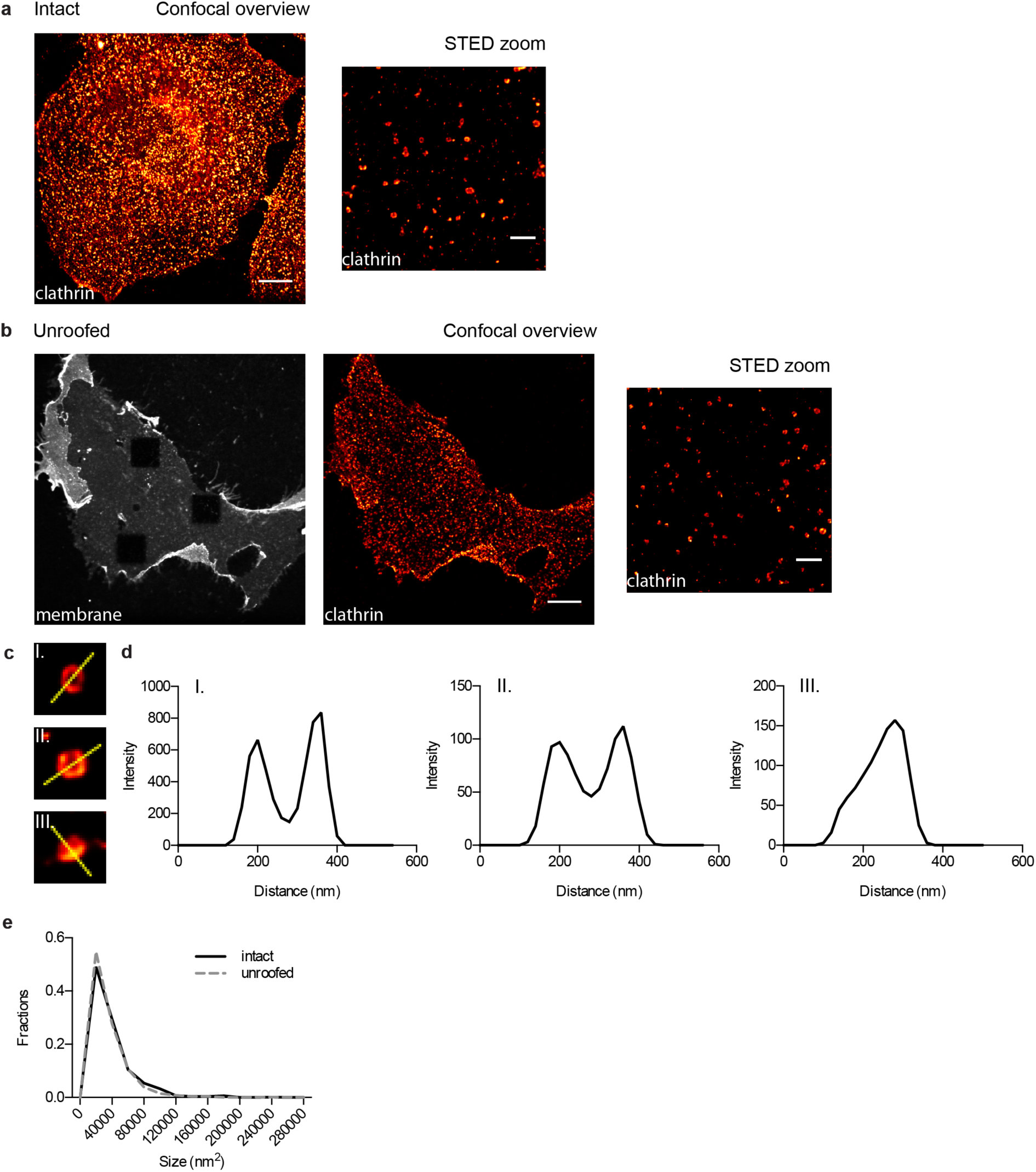
Comparative STED nanoscopy analysis of CCS in intact *vs.* unroofed cells. (a) STED nanoscopy of intact BSC-1 cells. CCS were immunostained with an anti-clathrin light chain (CLC) antibody. Left: confocal overview, scale bar: 5μm. Right: STED view, scale bar 1μm. (b) STED nanoscopy of unroofed BSC-1 cells. BSC-1 cells were unroofed by sonication; the remaining attached PM was stained with wheat germ agglutinin (WGA). CCS were immunostained with a an anti-clathrin antibody. Left: confocal overview, scale bar: 5μm. Right: STED view, scale bar 1μm. (c) Example of different CCS and their intensity line profile. Yellow line marks the axes measured for the intensity line profiles. (d) Analysis of the size distribution of CCS in intact and unroofed cells. CCS from five cells per conditions with three STED pictures per cell. Number of analysed CCS for intact cells: 820 and unroofed cells: 1169.

**Supplementary Figure 3:**
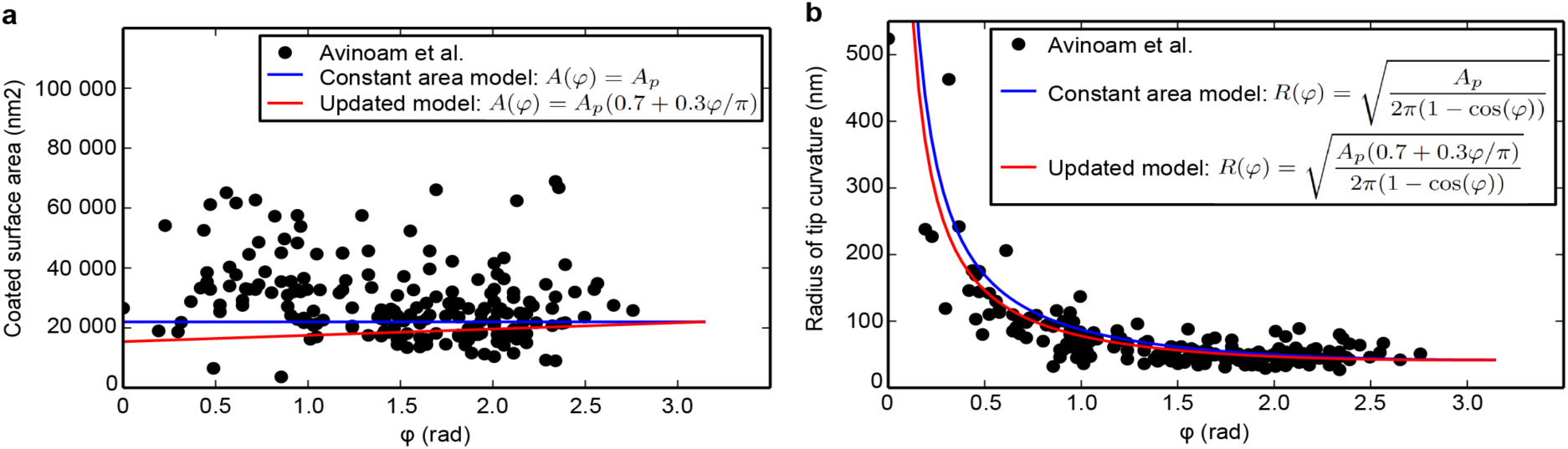
Updated growth model explains the CLEM data from Avinoam *et al.* (a) Clathrin coated surface area as a function of the growth angle *φ*. The constant area model where we assume A=22000nm^2^, corresponding to the most probable surface area^12^ is plotted (blue). Our updated model where we assume that the surface starts to bend when 70% of the final clathrin content is reached and grows the last 30% linear with *φ* plotted (red). The data (black dots) were extracted from Avinoam et al.^19^. (b) Radius of tip curvature as a function of the growth angle *φ*. Again, the constant area model (blue) and our updated model (red) are plotted. The radius of the tip curvature is given assuming that the clathrin structure exhibits the shape of a spherical cap. Then the tip curvature reads 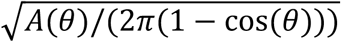. The data (black dots) were extracted from Avinoam et al.^19^.

## References

1. McMahon, H. T. & Boucrot, E. Molecular mechanism and physiological functions of clathrin-mediated endocytosis. Nat. Rev. Mol. Cell Biol. 12, 517–33 (2011).

2. Ferguson, S. M. & De Camilli, P. Dynamin, a membrane-remodelling GTPase. Nat Rev Mol Cell Biol 13, 75–88 (2012).

3. Taylor, M. J., Perrais, D. & Merrifield, C. J. A High Precision Survey of the Molecular Dynamics of Mammalian Clathrin-Mediated Endocytosis. PLoS Biol. 9, e1000604 (2011).

4. Heuser, J. Three-dimensional visualization of coated vesicle formation in fibroblasts. J. Cell Biol. 84, 560–583 (1980).

5. Maupin, P. & Pollard, T. D. Improved preservation and staining of HeLa cell actin filaments, clathrin-coated membranes, and other cytoplasmic structures by tannic acid-glutaraldehyde-saponin fixation. J. Cell Biol. 96, 51–62 (1983).

6. Larkin, J. M., Donzell, W. C. & Anderson, R. G. W. Potassium-dependent Assembly of Coated Pits: New Coated Pits Form as Planar Clathrin Lattices. J. Cell Biol. 103, 2619–2627 (1986).

7. Heuser, J. E. & Anderson, R. G. W. Hypertonic Media Inhibit Receptor-mediated Endocytosis by Blocking Clathrin-coated Pit Formation. J. Cell Biol. 108, 389–400 (1989).

8. Kirchhausen, T. Coated pits and coated vesicles — sorting it all out. Curr. Opin. Struct. Biol. 3, 182–188 (1993).

9. Kirchhausen, T. Imaging endocytic clathrin structures in living cells. Trends Cell Biol. 19, 596–605 (2009).

10. Saffarian, S., Cocucci, E. & Kirchhausen, T. Distinct dynamics of endocytic clathrin-coated pits and coated plaques. PLoS Biol. 7, e1000191 (2009).

11. den Otter, W. K. & Briels, W. J. The Generation of Curved Clathrin Coats from Flat Plaques. Traffic 12, 1407–1416 (2011).

12. Kumar, G. & Sain, A. Shape transitions during clathrin-induced endocytosis. Phys. Rev. E 94, 62404 (2016).

13. Lampe, M., Vassilopoulos, S. & Merrifield, C. Clathrin coated pits, plaques and adhesion. J. Struct. Biol. 196, 48–56 (2016).

14. Boucrot, E., Saffarian, S., Massol, R., Kirchhausen, T. & Ehrlich, M. Role of lipids and actin in the formation of clathrin-coated pits. Exp. Cell Res. 312, 4036–48 (2006).

15. Grove, J. et al. Flat clathrin lattices: stable features of the plasma membrane. Mol. Biol. Cell 25, 3581–3594 (2014).

16. Vassilopoulos, S. et al. Actin scaffolding by clathrin heavy chain is required for skeletal muscle sarcomere organization. J. Cell Biol. 205, 377–93 (2014).

17. Kirchhausen, T. & Harrison, S. C. Protein organization in clathrin trimers. Cell 23, 755–761 (1981).

18. Pearse, B. M. & Robinson, M. S. Purification and properties of 100-kd proteins from coated vesicles and their reconstitution with clathrin. EMBO J. 3, 1951–1957 (1984).

19. Avinoam, O., Schorb, M., Beese, C. J., Briggs, J. A. G. & Kaksonen, M. Endocytic sites mature by continuous bending and remodeling of the clathrin coat. Science 348, 1369 LP–1372 (2015).

20. Ehrlich, M. et al. Endocytosis by random initiation and stabilization of clathrin-coated pits. Cell 118, 591–605 (2004).

21. Sochacki, K. A., Shtengel, G., van Engelenburg, S. B., Hess, H. F. & Taraska, J. W. Correlative super-resolution fluorescence and metal-replica transmission electron microscopy. Nat Meth 11, 305–308 (2014).

22. Sochacki, K. A., Dickey, A. M., Strub, M.-P. & Taraska, J. W. Endocytic proteins are partitioned at the edge of the clathrin lattice in mammalian cells. Nat Cell Biol 19, 352–361 (2017).

23. Higgins, M. K. & McMahon, H. T. Snap-shots of clathrin-mediated endocytosis. Trends Biochem. Sci. 27, 257–263 (2017).

24. Wu, X. et al. Clathrin exchange during clathrin-mediated endocytosis. J. Cell Biol. 155, 291 LP–300 (2001).

25. Wu, X. et al. Adaptor and Clathrin Exchange at the Plasma Membrane and trans-Golgi Network. Mol. Biol. Cell 14, 516–528 (2003).

26. Saffarian, S. & Kirchhausen, T. Differential Evanescence Nanometry: Live-Cell Fluorescence Measurements with 10-nm Axial Resolution on the Plasma Membrane. Biophys. J. 94, 2333–2342 (2008).

27. Loerke, D., Mettlen, M., Schmid, S. L. & Danuser, G. Measuring the Hierarchy of Molecular Events During Clathrin-Mediated Endocytosis. Traffic 12, 815–825 (2011).

28. Boulant, S., Kural, C., Zeeh, J.-C., Ubelmann, F. & Kirchhausen, T. Actin dynamics counteract membrane tension during clathrin-mediated endocytosis. Nat Cell Biol 13, 1124–1131 (2011).

29. Ferguson, J. P. et al. Deciphering dynamics of clathrin-mediated endocytosis in a living organism. J. Cell Biol. 214, 347 LP–358 (2016).

30. Saleem, M. et al. A balance between membrane elasticity and polymerization energy sets the shape of spherical clathrin coats. Nat. Commun. 6, 6249 (2015).

31. Hassinger, J. E., Oster, G., Drubin, D. G. & Rangamani, P. Design principles for robust vesiculation in clathrin-mediated endocytosis. Proc. Natl. Acad. Sci. 114, E1118–E1127 (2017).

32. Robinson, M. S. Forty Years of Clathrin-coated Vesicles. Traffic 16, 1210–1238 (2015).

33. Maib, H., Smythe, E. & Ayscough, K. Forty years on: clathrin-coated pits continue to fascinate. Mol. Biol. Cell 28, 843–847 (2017).

34. Rao, Y. & Haucke, V. Membrane shaping by the Bin/amphiphysin/Rvs (BAR) domain protein superfamily. Cell. Mol. Life Sci. 68, 3983–3993 (2011).

35. Stachowiak, J. C. et al. Membrane bending by protein–protein crowding. Nat Cell Biol 14, 944–949 (2012).

36. Mim, C. & Unger, V. M. Membrane curvature and its generation by BAR proteins. Trends Biochem. Sci. 37, 526–533 (2017).

37. Ritter, B. et al. NECAP 1 regulates AP-2 interactions to control vesicle size, number, and cargo during clathrin-mediated endocytosis. PLoS Biol. 11, e1001670 (2013).

38. Miller, S. E. et al. CALM Regulates Clathrin-Coated Vesicle Size and Maturation by Directly Sensing and Driving Membrane Curvature. Dev. Cell 33, 163–175 (2015).

39. Merrifield, C. J., Perrais, D. & Zenisek, D. Coupling between clathrin-coated-pit invagination, cortactin recruitment, and membrane scission observed in live cells. Cell 121, 593–606 (2005).

40. Li, D. et al. Extended-resolution structured illumination imaging of endocytic and cytoskeletal dynamics. Science 349, (2015).

41. Heuser, J. E., Arnende, L. M., Branch, E. & Diseases, K. Deep-Etch Visualization of 27S Clathrin: A Tetrahedral Tetramer. J. Cell Biol. 105, 1999–2009 (1987).

42. Umasankar, P. K. et al. A clathrin coat assembly role for the muniscin protein central linker revealed by TALEN-mediated gene editing. Elife 3, e04137 (2014).

43. Cureton, D. K., Massol, R. H., Saffarian, S., Kirchhausen, T. L. & Whelan, S. P. J. Vesicular Stomatitis Virus Enters Cells through Vesicles Incompletely Coated with Clathrin That Depend upon Actin for Internalization. PLoS Pathog. 5, e1000394 (2009).

44. Godinez, W. J. & Rohr, K. Tracking Multiple Particles in Fluorescence Time-Lapse Microscopy Images via Probabilistic Data Association. IEEE Trans Med Imaging 34, 415–432 (2015).

